# Gain-of-function mutations in the RodZ juxtamembrane domain promote Rod system activity in *Escherichia coli*

**DOI:** 10.64898/2026.10.11.758364

**Authors:** Katherine A. Suarez, Thomas G. Bernhardt

## Abstract

The peptidoglycan (PG) cell wall protects bacteria from osmotic lysis and determines their shape. In rod-shaped bacteria, the lateral wall is elongated by the Rod system (elongasome). This synthetic machine is built around the RodA-PBP2 PG synthase complex, the actin-like protein MreB, and the accessory proteins MreC, MreD, and RodZ. Notably, RodZ shares a similar domain organization with FtsN, an essential division protein that triggers PG synthesis by the division apparatus (divisome). Both are single pass transmembrane proteins that bind actin-like proteins in the cytoplasm and have largely unstructured periplasmic regions capped by small, folded C-terminal domains. We therefore investigated whether RodZ might function like FtsN in activating the Rod system. To this end, gain-of-function variants of *Escherichia coli* RodZ were isolated that restored growth and shape to cells producing a hypomorphic *mreC* mutant. Production of these RodZ variants in wild-type cells increased resistance to the MreB antagonist A22 and enhanced sensitivity to the beta-lactam mecillinam. These phenotypes have all previously been associated with Rod system hyperactivation, and we found that they were caused by substitutions in the cytoplasmic juxtamembrane (JM) domain of RodZ. This domain is analogous to the N-terminus of FtsN known to promote divisome activation via its interaction with filaments of the actin-like FtsA protein. Structural modeling indicates that the JM domain of RodZ is similarly likely to associate with MreB filaments. Our results therefore support a model in which RodZ activates PG synthesis by the Rod system via a mechanism analogous to divisome activation by FtsN.

**IMPORTANCE:** Cell elongation in rod-shaped bacteria like *Escherichia coli* is mediated by the Rod system, and cell division is promoted by the divisome apparatus. It has long been recognized that these two machineries employ related components such as the peptidoglycan synthases RodA-PBP2 in the Rod system and FtsW-FtsI (PBP3) in the divisome. Here, we provide evidence that the RodZ protein of the Rod system plays a role in the activation of PG synthesis by the elongation machinery analogous to that performed by FtsN in divisome activation. The results suggest that the elongation and division machineries not only share related components but are also activated by similar regulatory mechanisms.

## INTRODUCTION

The peptidoglycan (PG) cell wall surrounds most bacterial cells and is essential for their integrity. In rod-shaped bacteria like *Escherichia coli*, two multiprotein machines build this structure: the Rod system that directs PG synthesis during cell elongation to promote rod-shape, and the divisome that forms the daughter cell poles by promoting PG synthesis at the division site (1) (**Fig 1**). Each system contains an essential PG synthase composed of a SEDS-family PG polymerase that synthesizes the glycan strands and a class B penicillin-binding protein (bPBP) that crosslinks them into the existing PG matrix (1–4). Both machineries also require an actin-like protein for their function (5, 6). Thus, the two systems are thought to share a common evolutionary origin (7). Mechanistic insights into the function and regulation of the Rod system have therefore facilitated advances in our understanding of the divisome and vice versa.

**Fig. 1.**
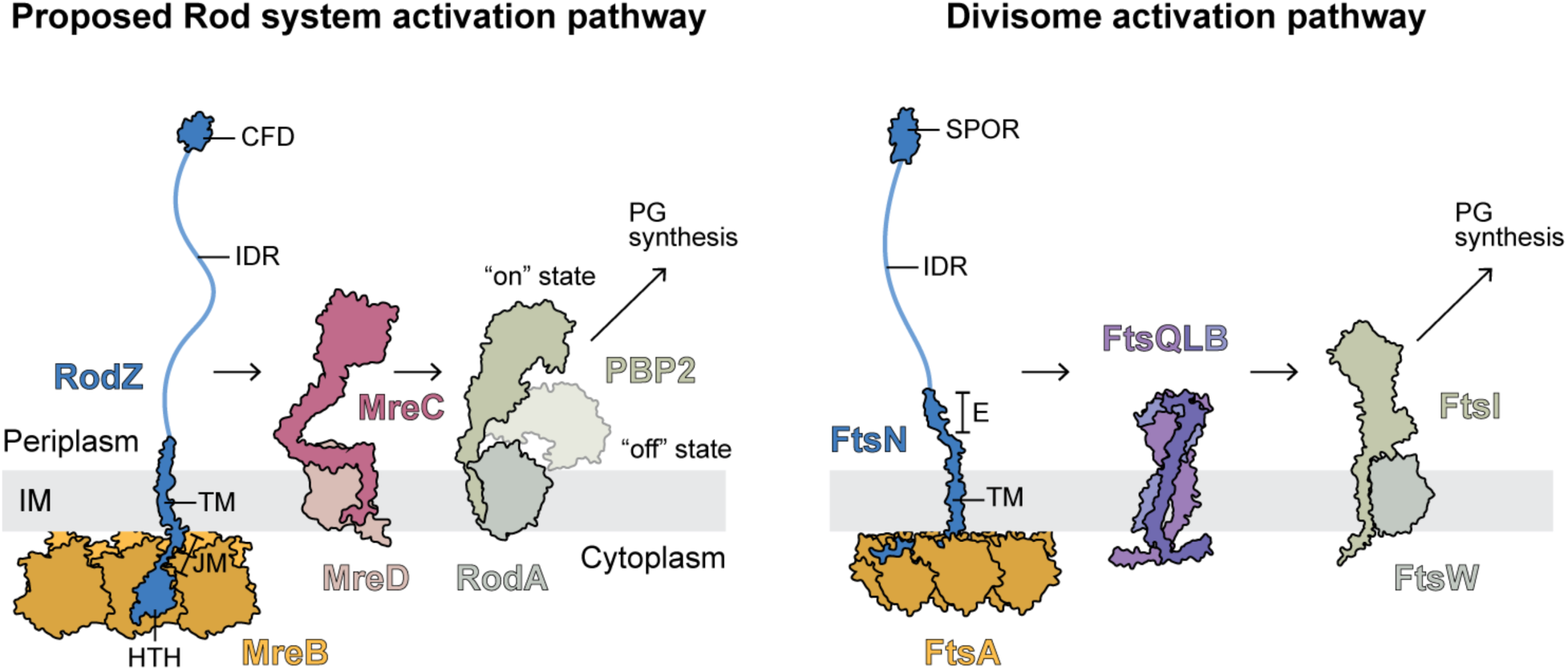
Models for the divisome and Rod system activation pathways. Schematic showing the components of the Rod system (**left**) and the divisome (**right**) with known of proposed roles in the activation of PG synthesis by RodA-PBP2 or FtsW-FtsI, respectively. See text for details.

A complex between RodA and PBP2 forms the essential SEDS-bPBP synthase of the Rod system (8, 9). Additional members of the system include MreB, MreC, MreD, and RodZ (1). MreB is the actin-like component (5). It forms twin, antiparallel filaments (10, 11) that are thought to serve as a molecular rudder to orient PG synthesis by the system orthogonal to the long cell axis (12). MreC, MreD, and RodZ are transmembrane proteins required for rod shape (13–16). RodZ binds to MreB via its cytoplasmic helix-turn-helix (HTH) domain (17), providing a connection between the membrane components and the MreB filaments. RodZ may also influence MreB polymerization (18). Genetic evidence suggests that MreC and MreD are regulatory components that control PG synthesis by the RodA-PBP2 complex (8, 19). Hypomorphic variants of MreC were shown to be suppressed by amino acid substitutions within RodA-PBP2 that activate its PG synthesis activity. Biophysical analysis using these and other engineered variants indicate that activation is associated with a conformational change in PBP2 that converts it from a membrane-proximal “down” state to a PG-proximal “up” state (20). These observations have led to a model in which the MreCD subcomplex stimulates a “down” to “up” conformational change in RodA-PBP2 to activate the Rod system (8, 20, 21) (**Fig 1**). How this activation is triggered has remained unclear.

Substitutions in the SEDS-bPBP synthase of the divisome called FtsW-FtsI (FtsWI) have also been shown to activate the division machinery (22–24). Because these substitutions map to locations analogous to those found to activate RodA-PBP2, the activation of FtsWI is thought to proceed via a similar conformational change in the complex. However, in the case of the divisome, activation is mediated by the FtsQ-FtsL-FtsB (FtsQLB) subcomplex, which is unrelated to MreCD (24–27) (**Fig 1**). The essential division protein FtsN has also been implicated in this activation mechanism (1). It is a multi-domain protein with: (i) a cytoplasmic N-terminus that interacts with the essential actin-like component of the divisome, FtsA (28), (ii) a transmembrane helix, (iii) an essential peptide positioned within a largely unstructured periplasmic domain (29, 30), and (iv) a C-terminal SPOR domain that binds a modified form of PG (31, 32) (**Fig 1**). FtsN is the last essential division protein to be recruited to the prospective division site before cell constriction is initiated (33). Its overproduction also suppresses the division defects of several different divisome mutants (34), and gain-of-function substitutions in the FtsQLB or FtsWI complex can bypass its essential function (23, 25, 26). The recruitment of FtsN to the division site is therefore thought to trigger a cascade that stimulates PG synthesis by FtsWI. It is proposed to do so via two different activation pathways: one using the periplasmic peptide that activates FtsWI via the FtsQLB complex and the other employing its cytoplasmic N-terminus and its interaction with the actin-like FtsA protein to stimulate PG synthesis by FtsWI from the cytoplasmic side of the membrane (35) (**Fig 1**).

Although they are not related by sequence, RodZ from the Rod system shares a common domain architecture with FtsN (**Fig 1**). Its cytoplasmic N-terminus includes an MreB-binding HTH domain and a positively charged juxtamembrane (JM) domain. These domains are followed by a single transmembrane (TM) helix and a periplasmic domain that like FtsN is also predicted to be largely disordered. Finally, this intrinsically disordered region (IDR) is capped by a small, C-terminal folded domain (CFD) of similar size to the SPOR domain. Additionally, activated variants of the Rod system PG synthase such as RodA-PBP2(L61R) can bypass RodZ function (8, 36) just as activated FtsWI variants render FtsN dispensable. Given the properties it shares with FtsN, we reasoned that RodZ might serve a triggering function for the Rod system analogous to FtsN in the divisome. To test this idea and define how RodZ contributes to Rod-system activation, we isolated gain-of-function variants of RodZ that suppress defects in the MreC component. These variants were found to have substitutions in the cytoplasmic JM region of RodZ, a domain previously shown to be critical for its function (14). The identified substitutions, Q98R and L102H, suppress the shape and viability defects of *mreC* hypomorphs and promote resistance to the MreB antagonist A22 (37). Although not sufficient for suppression, additional substitutions in the periplasmic domain of RodZ were found to enhance the suppressing activity of the Q98R substitution in the JM domain. Our genetic results therefore support a model in which RodZ activates the Rod system via a two-pronged activation mechanism analogous to divisome activation by FtsN (**Fig 1**). While this work was in progress, a related, independent study reported complementary results that also support this activation model (38).

## RESULTS

### Identification of RodZ variants that suppress mreC(R292H) defects

To identify hyperactive variants of RodZ, the *rodZ* coding sequence in plasmid pKS12 [P*_lac_*::*rodZ*] was mutagenized by error-prone PCR. The resulting plasmid library was transformed into strain PR5 harboring the *mreC*(R292H) mutation at the native chromosomal locus. The transformants were initially selected on minimal medium, conditions that are permissive for the growth of mutants defective for Rod system activity (39). They were then pooled and plated on the non-permissive condition, LB agar, in the presence of IPTG to induce the expression of *rodZ* from the plasmid. Cells harboring the empty vector gave rise to survivors on LB-IPTG agar at a frequency that was unaffected by induction, reflecting the background of chromosomal suppressors in the parent strain. Induction of the library raised the frequency of surviving colonies 31-fold above this background at 250 µM IPTG and 11-fold at 1 mM IPTG, indicating that most of the suppressors were likely to be plasmid encoded. To confirm that suppression was plasmid-linked, plasmids were purified from the survivors and transformed back into the PR5 [*mreC*(R292H)] parent strain. Plasmids that conferred suppression of the *mreC*(R292H) growth defect following this transformation were sequenced to map the location of the mutations responsible for the phenotype.

Three plasmid isolates (A13, B3, and B9) encoding *rodZ* suppressors were chosen for further study (**Table S1**). Isolates A13 and B9 each encoded RodZ with a Q98R substitution in the JM domain along with additional substitutions in either the CFD or IDR, respectively (**Table S1, Fig 2A**). Isolate B3 had a single substitution, L102H, in the JM domain (**Table S1, Fig 2A**). To identify the substitutions responsible for the phenotype, site-directed mutagenesis was used to generate constructs with individual changes to test for suppressing activity. Overproduction of RodZ(WT) failed to restore viability to the *mreC*(R292H) cells as expected (**Fig 2B**). By contrast, plasmids producing the Q98R and L102H variants of RodZ were sufficient for suppression of the *mreC*(R292H) growth defect (**Fig 2B**). Neither the A302V nor the substitutions in the IDR domain were sufficient for suppressing the *mreC*(R292H) growth defect (**Fig S1**). However, the combination of the three IDR substitutions (3IDR), but not the A302V change, enhanced the suppressing activity of the Q98R substitution (**Fig 2B and S1**). Additionally, production of a variant that combined the Q98R and L102H substitutions in the JM domain resulted in the best overall suppression of the *mreC*(R292H) growth defect with the suppression activity approaching that of the highly activated PBP2(L61R) variant isolated previously (8) (**Fig 2B**). The suppressors were not allele specific as they also restored the growth of cells harboring the *mreC*(G156D) hypomorphic allele (**Fig S2A**). Unlike the PBP2(L61R) variant, none of the RodZ variants suppressed the growth defect of Δ*mreCD* mutants (**Fig S2B**), indicating that residual MreCD activity is required for the RodZ variants to restore Rod system activity.

**Fig. 2.**
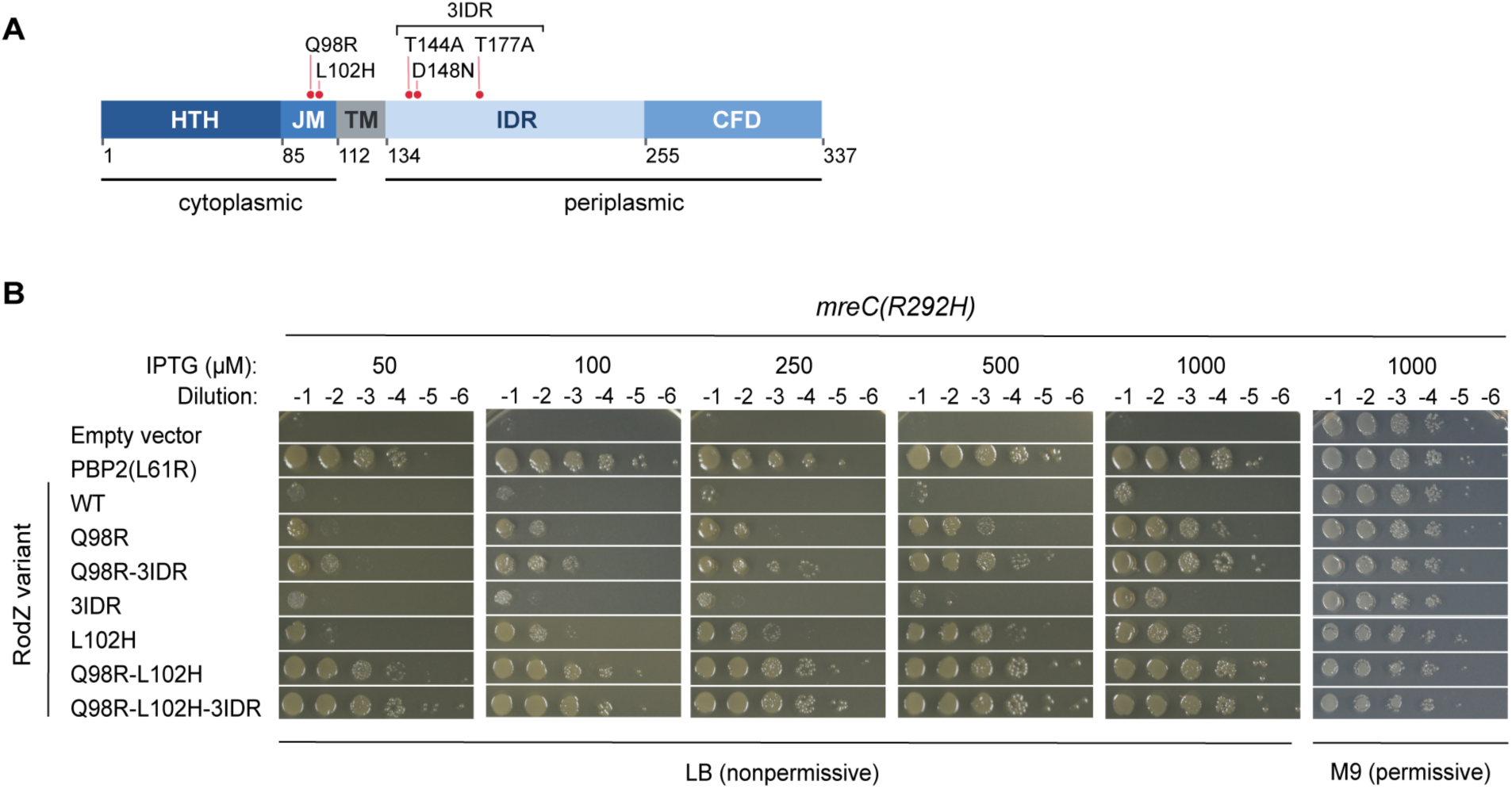
Substitutions in the JM domain of RodZ suppress the growth defect of an *mreC*(R292H) mutant. **(A)** Domain map of RodZ. Full-length RodZ (337 residues) comprises a cytoplasmic helix-turn-helix domain (HTH, residues 1-84) and juxtamembrane region (JM, 85-111), a single transmembrane helix (TM, 112-133), a periplasmic intrinsically disordered region (IDR, 134-254), and a C-terminal folded domain (CFD, 255-337). Amino acid substitutions are indicated as red-filled circles. **(B)** Cells of PR5 *mreC*(R292H) carrying the indicated P_lac_ expression plasmids, pMT77 (empty vector), pPR122 (PBP2(L61R)), pKS12 (RodZ), pKS49 (RodZ(Q98R)), pKS91 (RodZ(Q98R-3IDR)), pKS90 (RodZ(3IDR)), pKS53 (RodZ(L102H)), pKS61 (RodZ(Q98R-L102H)), and pKS92 (RodZ(Q98R-L102H-3IDR)), were serially diluted, and spotted as described in Materials and Methods.

We next tested the ability of the RodZ variants to suppress the shape defect of *mreC*(R292H) cells. The results largely mirrored the growth assays. The variants with single substitutions in the JM domain, RodZ(Q98R) and RodZ(L102H), each resulted in a partial shape restoration as assessed by measuring cell width (**Fig 3**). Again, the combination of these substitutions led to the most robust suppression with the width of the cells approaching that of the *mreC*(R292H) mutant producing the hyperactive PBP2(L61R) variant (**Fig 3**). As with the growth assays, combining the 3IDR substitutions with the Q98R change enhanced suppression activity (**Fig 3**). Notably, the 3IDR substitution variant resulted in mild suppression of the *mreC*(R292H) shape defect even without the additional JM substitution (**Fig 3**), suggesting that the 3IDR substitutions alone have suppression activity. Immunoblot analysis indicated that all the variants were produced at levels roughly equivalent to RodZ(WT) (**Fig S3**). We therefore conclude that changes in the JM domain of RodZ and, to a lesser degree, the periplasmic IDR region enhance RodZ activity.

**Fig. 3.**
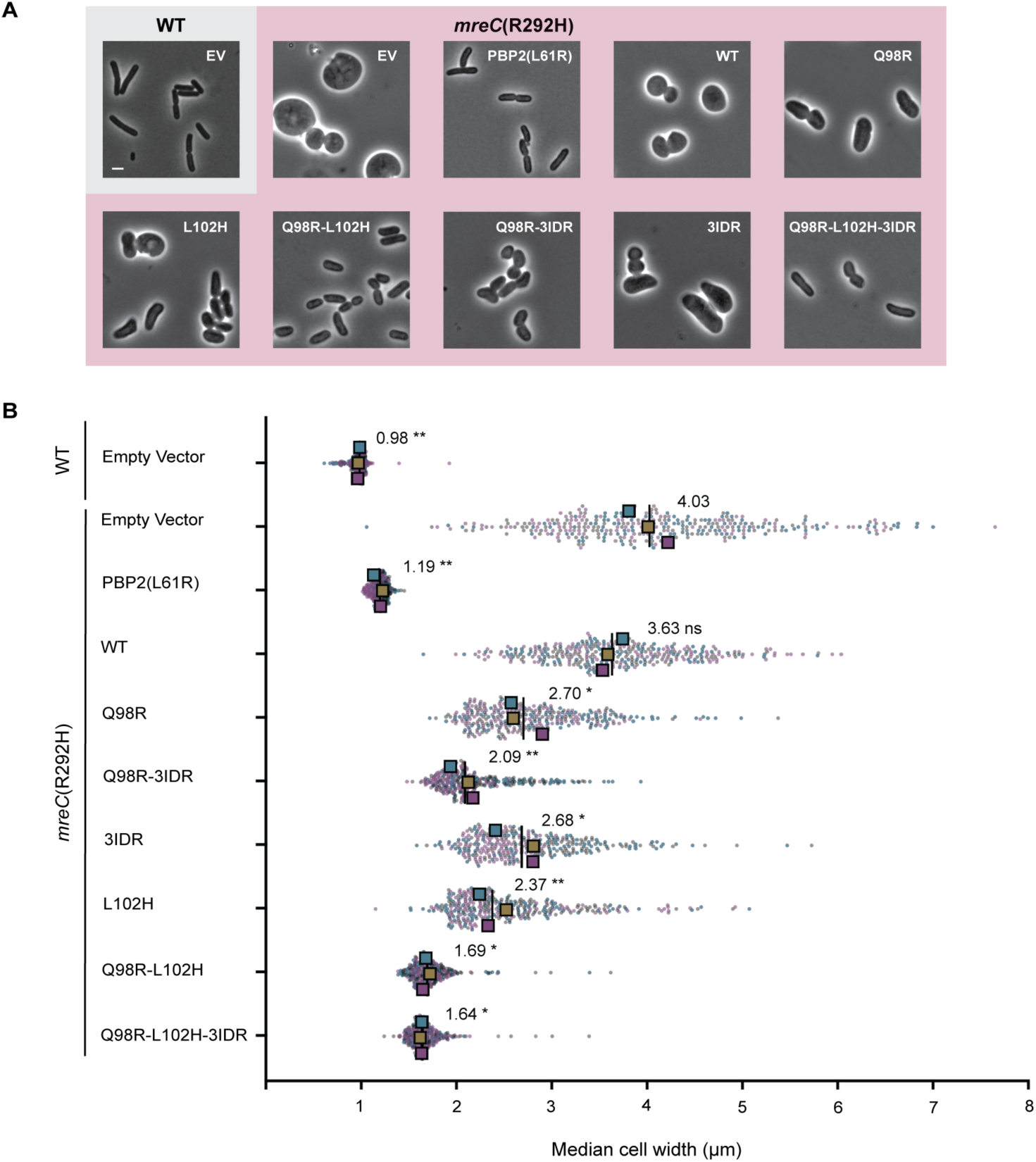
RodZ variants suppress the shape defect of *mreC*(R292H) cells. **(A)** Representative phase-contrast micrographs of WT (MG1655/pMT77) and *mreC*(R292H) (PR5) cells carrying the plasmids listed in Fig 2, grown and imaged as described in Materials and Methods. Scale bar, 2 µm. **(B)** The median width of each cell was quantified as described in Materials and Methods. Small points show single-cell measurements, colored by biological replicate. Large squares show the median of each replicate population (n = 3 independent transformations, 118 cells per replicate), colored to match their replicate. Vertical lines and printed values show the mean of the three replicate medians. Statistical testing was performed on replicate medians by Brown-Forsythe and Welch ANOVA with Dunnett’s T3 multiple-comparison correction; asterisks indicate comparison with *mreC*(R292H) carrying the empty vector. *, P ≤ 0.05; **, P ≤ 0.01; ***, P ≤ 0.001; ****, P ≤ 0.0001; ns, not significant.

### Evidence that the RodZ variants hyperactivate the Rod system

Variants of the RodA-PBP2 synthase that overcome Rod system defects hyperactivate the Rod system when produced in the context of an otherwise wild-type background (8). Hyperactivation of the Rod system results in increased resistance to the MreB antagonist A22 and hypersensitivity to the beta-lactam mecillinam, which induces a toxic malfunction of the system (40). Therefore, to determine if the RodZ variants with JM domain substitutions also hyperactivate the Rod system, we monitored their effect on the A22 resistance and mecillinam sensitivity of wild-type cells. Production of RodZ variants with individual Q98R or L102H substitutions led to a modest increase in A22 resistance whereas the RodZ(Q98R-L102H) variant with both substitutions promoted a robust increase in A22 resistance, mirroring that induced by PBP2(L61R) production (**Fig 4A**). Similarly, the RodZ(Q98R-L102H) variant caused an increase in mecillinam sensitivity that was comparable to that induced by PBP2(L61R) (**Fig 4B**). Based on these results, we conclude that the JM domain substitutions in RodZ promote Rod system activity.

**Fig. 4.**
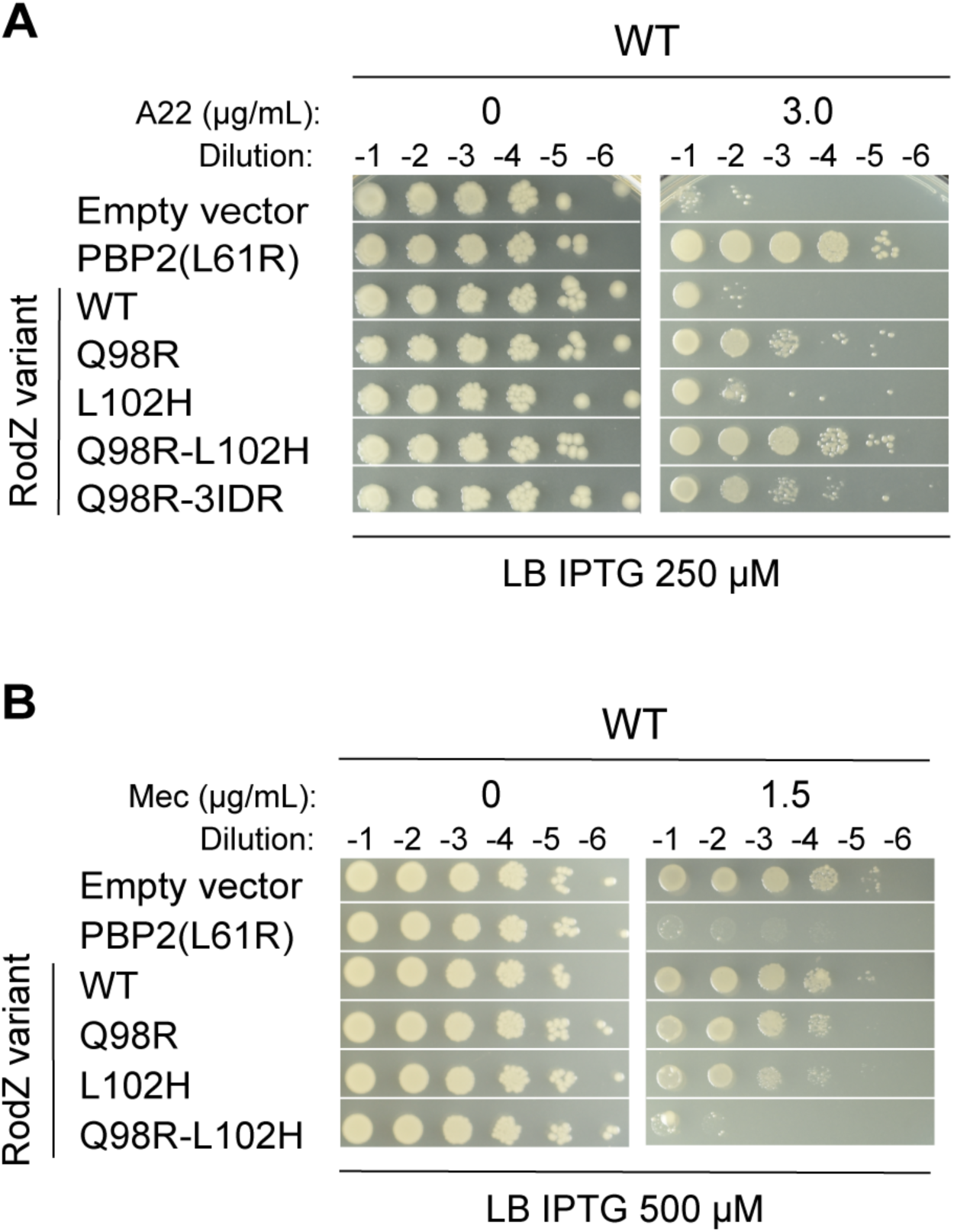
RodZ variants promote resistance to A22 and increased sensitivity to mecillinam. MG1655 [WT] cells carrying the indicated P_lac_-expression plasmids were grown, serially diluted, and spotted (5 µl) onto LB CM25 plates containing the indicated IPTG and drug concentrations. Plates were incubated for 18 h at 37°C before imaging. **(A)** A22 challenge. Plates contained 250 µM IPTG and A22 at 0 or 3.0 µg/mL. Plasmids: pMT77 (empty vector), pPR122 (PBP2(L61R)), pKS12 (RodZ), pKS49 (RodZ(Q98R)), pKS53 (RodZ(L102H)), pKS61 (RodZ(Q98R-L102H)), pKS91 (RodZ(Q98R-3IDR)), and pKS92 (RodZ(Q98R-L102H-3IDR)). **(B)** Mecillinam challenge. Plates contained 500 µM IPTG and mecillinam at 0 or 1.5 µg/mL. Plasmids: pMT77 (empty vector), pPR122 (PBP2(L61R)), pKS12 (RodZ), pKS49 (RodZ(Q98R)), pKS53 (RodZ(L102H)), and pKS61 (RodZ(Q98R-L102H)).

### Domain requirements for the suppressing activity of RodZ variants

To identify the regions of RodZ required for the changes in the JM domain to overcome Rod system defects, a series of truncation variants were constructed and tested for their ability to suppress the *mreC*(R292H) or *mreC*(G156D) growth defects. Truncation of the periplasmic domain at residue 254 or 155 did not appreciably reduce the suppressing activity promoted by the combined Q98R and L102H substitutions in the JM domain (**Fig 5A-B**). The same truncations modestly reduced the activity of the single JM substitution variants (**Fig 5A-B**). To test the requirement for MreB binding by the HTH domain, we initially constructed an N-terminal deletion that removed this region of the protein. However, immunoblot analysis indicated that this truncated protein was unstable and failed to accumulate (**Fig S3**). We therefore constructed variants with an F60A substitution previously shown to disrupt the interface between the HTH domain and MreB (17). The F60A change abolished suppression by the single Q98R and L102H substitution variants but was tolerated by the Q98R-L102H double substitution variant, which retained strong suppression activity (**Fig 5C and S4**). Notably, the F60A change also severely attenuated the suppressing activity of the RodZ(Q98R-3IDR) variant despite its original suppressing activity being nearly as good as the RodZ(Q98R-L102H) variant. Thus, bypass of the HTH-MreB contact depends on the double substitution in the JM domain and is not simply a function of the overall suppression strength of a variant. Immunoblot analysis indicated that all variants except for the HTH truncation accumulate to levels equivalent to RodZ(WT) (**Fig S5**). We conclude that the RodZ(Q98R-L102H) variant can promote Rod system activity independent of MreB binding by the HTH domain.

**Fig. 5.**
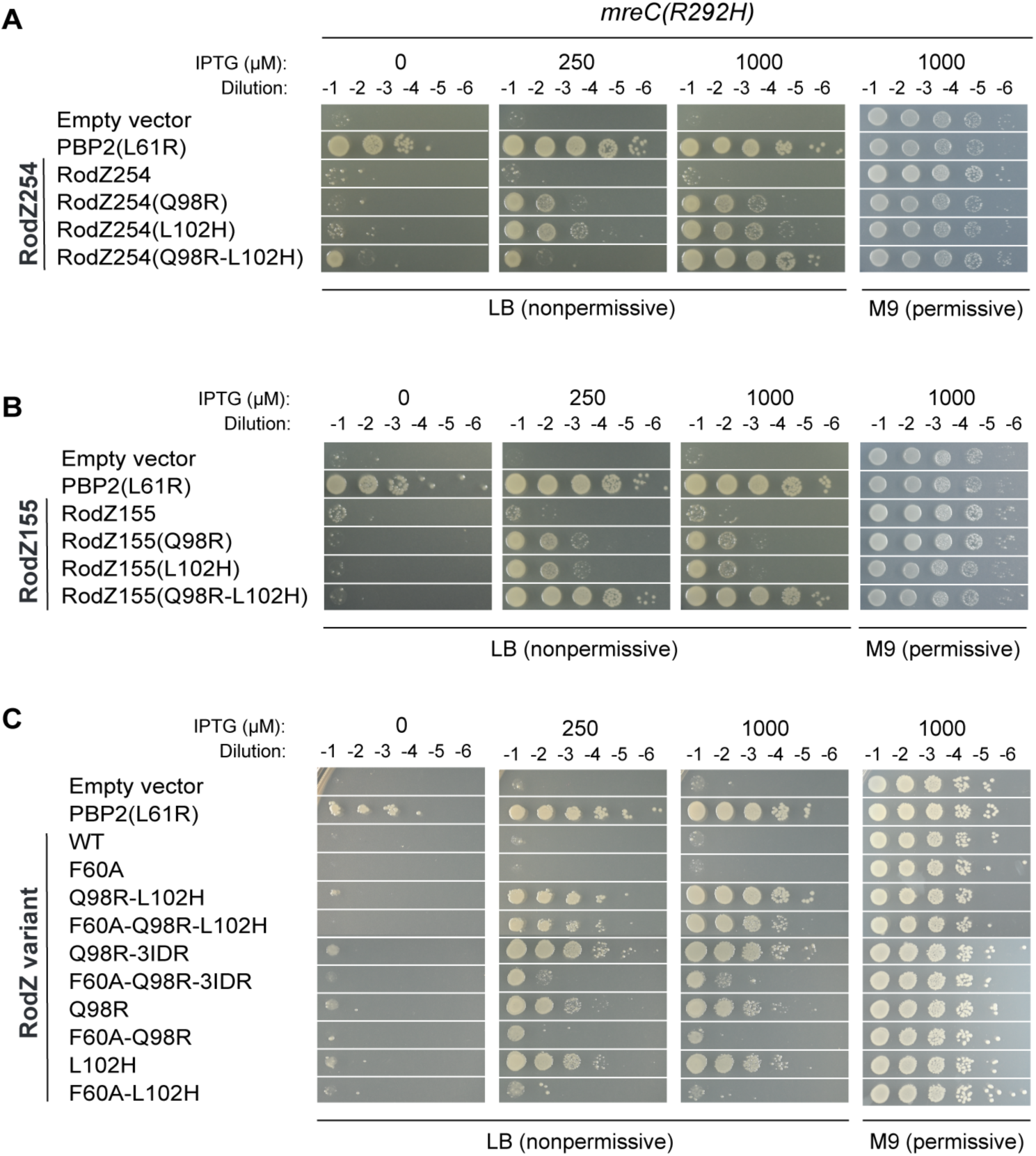
Domain requirements for RodZ variants to suppress *mreC*(R292H). Cells of PR5 [*mreC*(R292H)] carrying the indicated P_lac_ expression plasmids were grown, normalized, and spotted on the indicated media as described in Materials and Methods. **(A)** <u>RodZ</u> (1-254) <u>set</u>: pMT77 (empty vector), pKS28 (RodZ254), pPR122 (PBP2(L61R)), pKS66 (RodZ254(Q98R)), pKS67 (RodZ254(L102H)), and pKS68 (RodZ254(Q98R-L102H)). **(B)** <u>RodZ</u> (1-155) <u>set</u>: pMT77 (empty vector), pKS13 (RodZ155), pPR122 (PBP2(L61R)), pKS63 (RodZ155(Q98R)), pKS64 (RodZ155(L102H)), and pKS65 (RodZ155(Q98R-L102H)). **(C)** <u>RodZ(F60A) set</u>: pMT77 (empty vector), pPR122 (PBP2(L61R)), pKS12 (RodZ), pKS116 (RodZ(F60A)), pKS117 (RodZ(F60A-Q98R-L102H)), pKS118 (RodZ(F60A-Q98R-3IDR)), pKS119 (RodZ(F60A-Q98R)), and pKS120 (RodZ(F60A-L102H)).

## DISCUSSION

RodZ has long been known to be a critical component of the Rod system, but our understanding of its function remains incomplete. Most of what we know about the role of RodZ within the Rod system is derived from protein-protein interaction studies and loss-of-function mutants encoding truncated RodZ variants or deletion mutants that lack the protein entirely. In cells inactivated for RodZ, MreB localization was shown to be altered such that its distribution is more patchy and less uniform (14). These cells have also been found to assemble fewer MreB polymers (41). It has been well established that RodZ interacts with MreB via its HTH domain (17). Two-hybrid and other assays also suggest that RodZ is also likely to directly or indirectly associate with the transmembrane proteins of the Rod system (14). Based on these results, RodZ is commonly thought to have two functions: one involved in modulating MreB filament formation or stability and the other in linking MreB filaments to the membrane and other components of the Rod system.

To gain additional insight into RodZ function, we selected for hyperactive RodZ variants capable of suppressing the growth and shape defects of cells encoding a hypomorphic derivative of MreC. The selection identified RodZ variants with changes in the JM domain (Q98R and L102H) that are not only capable of suppressing MreC defects but also promote other hallmarks of Rod system hyperactivation such as enhanced A22 resistance and mecillinam hypersensitivity. It is not currently feasible to reconstitute the full Rod system in a purified system to test the effect of these altered RodZ variants on Rod system activity directly. However, given the similar phenotypes induced by the production of RodZ(Q98R-L102H) with those promoted by PBP2(L61R), a variant known to activate PG synthesis by RodA-PBP2 *in vitro* (8), a reasonable explanation for the effect of the JM domain substitutions in RodZ is that they too act by directly or indirectly stimulating PG biogenesis by RodA-PBP2. Thus, our results suggest that RodZ is more than a mere linker connecting MreB filaments to the RodA-PBP2 synthase and other membrane components of the Rod system but that it also may form part of the regulatory cascade that activates PG synthesis by the complex. Complementary results studying hyperactive MreC variants and the requirement of RodZ domains for Rod system function were also recently reported that support a role for RodZ in Rod system activation (38).

### Possible mechanism by which the JM substitutions promote Rod system activity

The RodZ(Q98R-L102H) variant promotes Rod system activity even when its interaction with MreB via the HTH domain is disrupted. It is therefore possible that the JM domain also provides additional contacts between RodZ and MreB. Indeed, prior truncation studies found the JM domain to be the most critical for RodZ function (14), and bimolecular fluorescence complementation assays also indicate that RodZ lacking its HTH domain remains capable of interacting with MreB (42). A JM-MreB contact is also supported by AlphaFold modeling (43) of the RodZ HTH and JM domain (residues 1-111) structure in complex with an MreB filament (**Fig 6**). The model predicts that the MreB subunits form a short segment of a paired antiparallel filament. These paired filaments have previously been implicated as the functional form of the MreB rudder that engages with the Rod system and orients PG synthesis (11, 12). As expected from the known structure (17), the model predicts a contact between the HTH domain of RodZ and one of the MreB monomers. However, the model also positions the JM domain near the interface between two MreB monomers of one of the filaments (**Fig 6**). Importantly, both the Q98 and L102 residues map to this predicted interface between RodZ and MreB (**Fig 6**), suggesting that it is biologically relevant. Thus, we propose that the JM domain of RodZ interacts with MreB and that the Q98R and L102H substitutions enhance the effect of the interaction on Rod system function to overcome defects caused by substitutions in MreC.

**Figure 6.**
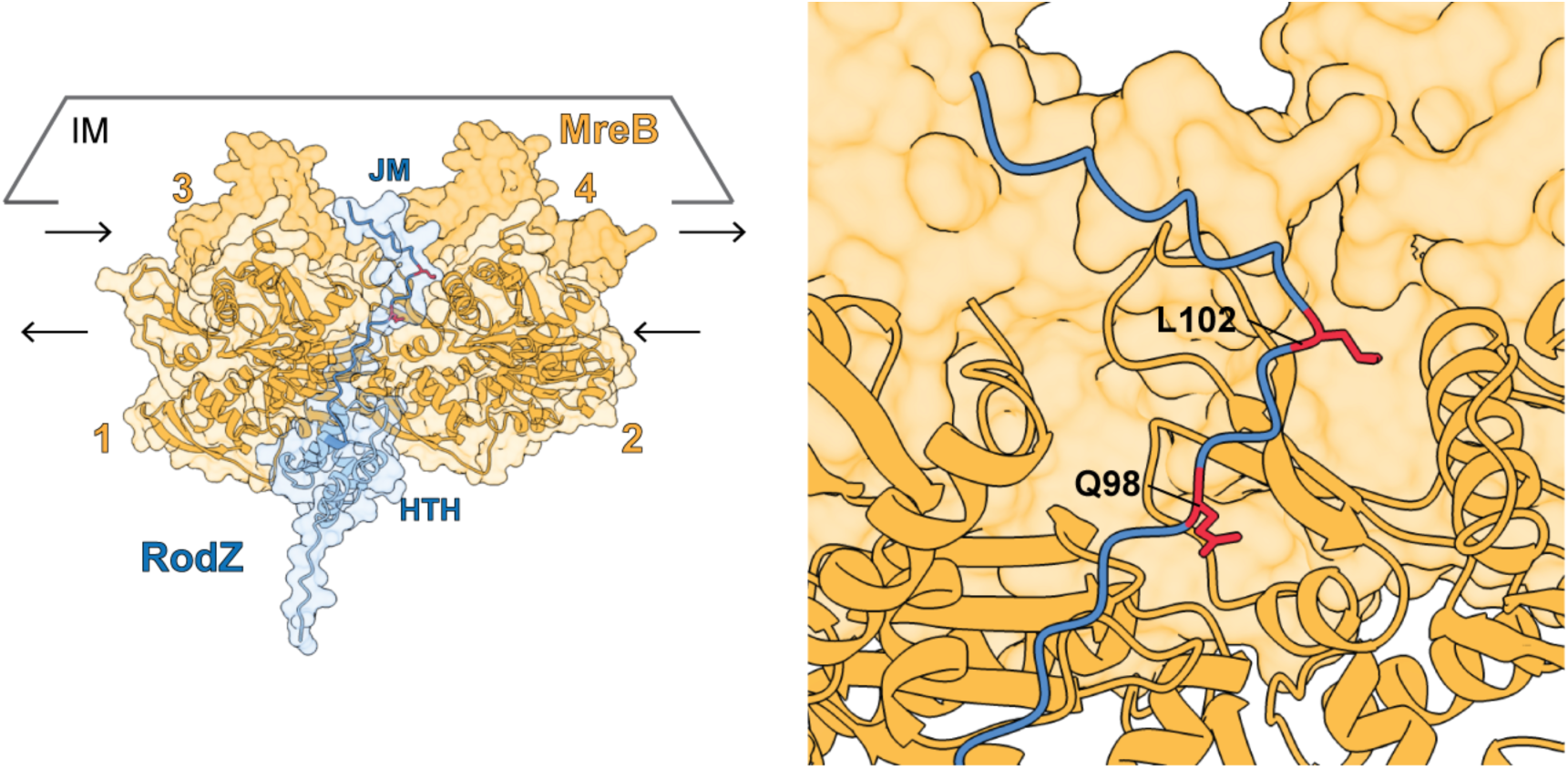
The JM domain of RodZ is predicted to interact with MreB. **(A)** AlphaFold3 model of the RodZ helix-turn-helix (HTH) and juxtamembrane (JM) regions (residues 1-111) folded with six subunits of MreB. Only four MreB subunits are shown. RodZ HTH, light blue; RodZ JM, dark blue; MreB, gold. Black arrows indicate the orientation of the two antiparallel MreB filaments. The plane of the inner membrane (IM) is outlined in gray. **(Right)** Close-up of the predicted JM-MreB binding site highlighting the residues altered in the hyperactive RodZ variants. Confidence metrics of the model were as follows: global ipTM = 0.74, HTH-MreB contact ipTM = 0.51 and pLDDT = 70.7, JM-MreB contact ipTM = 0.34 and pLDDT = 40.7.

### Role of the periplasmic domain?

The periplasmic IDR domain of RodZ is predicted to be largely unstructured. If the JM domain is intact, this domain can be deleted without severely compromising RodZ function (14). Thus, it is possible that the periplasmic domain of RodZ plays a role in Rod system function that is partially redundant with the JM domain. Consistent with this possibility, the three substitutions in the IDR region identified in the suppressor analysis were found to enhance RodZ activity, albeit weakly compared to the changes in the JM domain. We therefore propose that the periplasmic IDR domain of RodZ may also promote Rod system activity, potentially via an interaction with the periplasmic domains of other components of the complex. In support of this model, a recent report has also implicated this region of RodZ in Rod system activation and evidence is presented that it does so via an interaction with MreC (38).

In addition to the IDR domain, the periplasmic region of RodZ also has a small, folded domain of unknown function at its C-terminus. When the predicted structure of this CFD was used to perform a structure-based similarity search on the DALI server (44), the search revealed moderate similarity with the family 60 carbohydrate-binding module vCBM60 (PDB 2XHH, Z-score = 5.8)

(45). The canonical CBM60 module recognizes carbohydrates through a defined interaction involving an aromatic platform that stacks against the sugar rings, a coordinating calcium ion, and an open cleft formed by two loops. The RodZ CFD does not contain these features at the analogous location of its predicted structure, suggesting that if it binds a carbohydrate, it may do so in a manner distinct from the CBM60 module. Nevertheless, the finding that this domain of RodZ shares similarity with a carbohydrate-binding domain raises the intriguing possibility that it might function analogously to the SPOR domain of FtsN by binding PG and modulating RodZ localization.

### Additional parallels with FtsN and divisome activation

The identification of substitutions in the JM and periplasmic IDR domains of RodZ that promote Rod system function lends further support to the idea that RodZ and FtsN are functional analogs. The N-terminal juxtamembrane domain of FtsN has been shown to interact with the actin-like FtsA protein (28). This interaction has been shown to promote the formation of paired antiparallel FtsA filaments, and this change in FtsA polymer structure has been proposed to be part of the divisome activation pathway (46). A similar transition into an antiparallel filament structure for MreB may therefore also be involved in Rod system activation. Furthermore, the hyperactivity caused by substitutions in the RodZ JM domain and AlphaFold modeling suggest that an interaction between the JM domain of RodZ and MreB may stimulate this change. In support of this possibility, the MreB(A125V) variant was previously shown to suppress the growth and shape phenotype of a Δ*rodZ* mutant (36). This substitution maps to the interface between antiparallel filaments of MreB (11), suggesting the A125V change may stabilize the paired filament structure to bypass the need for RodZ. We therefore propose that the JM domain of RodZ functions to provide a cytoplasmic signal for Rod system activation via its effect on MreB filament structure, analogous to the role of the N-terminal juxtamembrane domain of FtsN in modulating FtsA filament structure to promote divisome activation.

Like RodZ, the periplasmic domain of FtsN is largely unstructured. The essential domain (peptide) of FtsN is embedded within this unstructured region and has been implicated in providing a periplasmic signal for divisome activation via the FtsQLB complex (25, 26, 30). Although further work will be required to investigate this possibility, the substitutions in the periplasmic IDR of RodZ that promote Rod system activity may similarly identify a periplasmic activation signal provided by RodZ that, in turn, may act via the MreC and/or MreD components, as suggested by recent results (38). Thus, the Rod system, like the divisome, may utilize a combination of cytoplasmic and periplasmic signals provided by RodZ to ensure the complex is fully assembled before PG synthesis is activated.

## MATERIALS AND METHODS

### Media, bacterial strains, and plasmids

All *Escherichia coli* strains used in this study are derivatives of MG1655 (47) and are listed in **Table S2**. Strains were grown in LB (1% tryptone, 0.5% yeast extract, 0.5% NaCl) or minimal M9 medium supplemented with 0.2% casamino acids and 0.2% glucose (abbreviated M9 CAA glu). Rod system mutants were maintained on M9 CAA glu at 30°C unless otherwise indicated, as the Rod system is dispensable under this condition and essential on rich medium (39). Unless otherwise indicated, chloramphenicol was used at 25 µg/mL (CM25) and all growth was at 30°C except for the A22 and mecillinam assays, which were performed at 37°C. Growth conditions for individual experiments are described in the figure legends.

All plasmids are listed in **Table S3** and all primers in **Table S4**. Detailed information about plasmid construction can be found in Supplemental Material. Substitutions were introduced into pKS12 [P_lac_::*rodZ*] and its derivatives by whole-plasmid PCR with Q5 High-Fidelity polymerase (New England Biolabs) followed by KLD treatment (New England Biolabs, M0554) and transformation into chemically competent cells. All plasmids were sequence verified by the DNA Resource Core of Dana-Farber/Harvard Cancer Center.

### Construction of the mutagenized *rodZ* plasmid library

A library of *rodZ* alleles was constructed in pKS12, a p15A-origin, chloramphenicol-resistant vector carrying the full-length 1,011-bp *rodZ* coding sequence under P_lac_ control, using a megaprimer-based whole-plasmid mutagenesis strategy adapted from the GeneMorph II EZClone Domain Mutagenesis Kit protocol (Agilent, catalog #200552). Mutagenized megaprimers spanning *rodZ* and approximately 150 bp of flanking sequence on each side (1.3 kb total) were generated in five independent low-fidelity PCR reactions (pools 1-5) using *Taq* DNA polymerase, primers #76 and #96, and drop-dialyzed pKS12 template. An extension temperature of 68°C rather than 72°C was used, as the higher temperature generated a spurious secondary product. Each purified megaprimer pool (500 ng) was then used to amplify the full plasmid with Q5 High-Fidelity master mix, and parental template was removed by DpnI digestion.

DpnI-treated products were drop-dialyzed and electroporated into NEB 10-beta cells (New England Biolabs), and the resulting library DNA was re-transformed into electrocompetent PR5. Each pool yielded in excess of 10⁶ transformants and the no-DNA controls yielded no colonies.

Colonies were harvested by scraping, adjusted to OD_600_ = 10, and stored at −80°C in LB containing 10% glycerol or 10% DMSO. To estimate mutation frequency, the mutagenized region was Sanger sequenced from randomly selected transformants. Of the nine clones tested, eight (88%) carried at least one substitution within the amplified region, with substitutions distributed across the *rodZ* coding sequence and an average of approximately one substitution per allele.

### Selection for suppressors of *mreC*(R292H)

The five library pools, together with a pool combining all five, were transformed into PR5 [*mreC*(R292H)], with pMT77 (empty vector), pKS12, and pPR122 [P_lac_::*pbpA*(L61R)] transformed in parallel as negative and positive controls. Overnight cultures of the resulting transformants were serially diluted and plated on permissive medium (M9 CAA glu CM25) or non-permissive medium (LB CM25), each containing IPTG at 0, 250 µM, or 1 mM, and incubated at 30°C for 40 h (M9 CAA glu) or 18 h (LB).

Fifty colonies were picked from large-format (150 mm) LB plates, 25 each from the 250 µM and 1 mM IPTG conditions, and streak-purified on both permissive and non-permissive media at all three IPTG concentrations. Plasmids were isolated from these isolates and re-transformed into fresh PR5 alongside the controls above, and isolates showing IPTG-dependent growth under non-permissive conditions were re-tested two to three additional times to confirm that the phenotype segregated with the plasmid rather than with a chromosomal change. Three isolates were validated by this criterion and the plasmids were sequenced to map the responsible mutations (**Table S1**).

### Viability assays

Strains carrying the indicated P_lac_-driven plasmids were grown at 30°C, normalized to equal cell density (OD_600_ = 1), serially diluted tenfold, and spotted (5 µl) onto M9 CAA glu CM25 (permissive) and LB CM25 (non-permissive) containing IPTG across the range indicated for each figure. LB plates were incubated for 18 h and M9 CAA glu plates for 40 h at 30°C, unless otherwise noted. Dilution factors are indicated above the spot series in each figure.

### Image acquisition and analysis

Cultures were grown overnight in M9 CAA glu CM25 without IPTG at 30°C, back-diluted to OD_600_ = 0.05 in M9 CAA glu CM25 with 250 µM IPTG and grown to OD_600_ ≈ 0.2. Cells were then diluted to an OD_600_ = 0.05 in LB (no CM25) with 250 µM IPTG and grown to an OD_600_ ≈ 0.2 before being fixed in 2.6% formaldehyde with 0.04% glutaraldehyde at room temperature for 1 h. Prior to imaging, cells were pelleted (8,000 × *g*), concentrated 10×, immobilized on 1.7% agarose pads containing M9 CAA glu, and covered with #1.5 coverslips.

Phase-contrast microscopy was performed on a Nikon TE2000 inverted microscope equipped with a Plan Apo 100×/1.45 NA Oil Ph3 objective. Images were acquired using Nikon Elements 5.2 acquisition software Cropping and display adjustments were performed in Fiji (48). Median cell widths were measured using the Fiji plugin MicrobeJ (49) from three independent replicate populations of 118 cells each (*n* = 354 cells per strain) and displayed as SuperPlots (50). For each strain, three biological replicates were analyzed, each derived from an independent transformation and processed on separate days.

### Immunoblotting

Whole-cell extracts were prepared from cultures grown under the conditions described for phase-contrast microscopy. Protein concentrations were determined using a copper-based colorimetric assay (G-Biosciences NI Protein Assay) against a BSA standard curve. Samples were normalized, loaded at 20 µg of total protein per lane, run on 4-20% gradient polyacrylamide gels, and transferred to an activated PVDF membrane using a semi-dry transfer system (Bio-Rad Trans-Blot Turbo). Membranes were blocked in 5% nonfat milk in PBS-T (PBS, 0.1% Tween-20) for 1.5 h at room temperature, then transferred to primary antibody solution containing rabbit polyclonal antiserum raised against *E. coli* RodZ(1–319) (rabbit 01806; (14)) at 1:15,000 in 1% milk-PBS-T and incubated for 16 h at 4°C. After four washes in PBS-T, membranes were incubated with anti-rabbit IgG HRP secondary antibody (Rockland TrueBlot, cat# 18-8816-33) at 1:5,000 in 0.2% milk-PBS-T for 2 h at room temperature, followed by five washes in PBS-T. Blots were developed using SuperSignal West Pico PLUS Chemiluminescent Substrate (Thermo Fisher, cat# 34580) and imaged on a Bio-Rad ChemiDoc MP Imaging System. Membranes were also stained with Ponceau S to confirm equal loading. Full-length RodZ has a predicted molecular weight of 37 kDa but migrates anomalously at 55 kDa, consistent with previous reports (14, 15).

### Structural modeling and analysis

Sequences for RodZ and MreB correspond to *E. coli* K-12 substr. MG1655 (GenBank U00096.3; annotation derived from EcoCyc). Except where noted, structural models were generated using the AlphaFold Server (AlphaFold3) (43) with default server settings. RodZ(1-111), comprising the HTH and JM regions, was modeled with a six-subunit MreB filament (only four are shown in the figure). Contacts between RodZ residues Q98 and L102 and MreB were scored for reproducibility across the 5 models per job and across occupancy conditions. Confidence metrics of the model were as follows: global ipTM = 0.74, HTH-MreB contact ipTM = 0.51 and pLDDT = 70.7, JM-MreB contact ipTM = 0.34 and pLDDT = 40.7.

The top-ranked of five AlphaFold3 models of the RodZ CFD (residues 255-337) was submitted as a single chain to the DALI server (44) for comparison against the PDB25 representative set (PDB chains sharing < 25% pairwise sequence identity) in June 2026. Following established guidance, matches with Z < 2 were treated as non-significant and those with Z-scores of 2-8 as fold-level resemblances requiring further evaluation rather than evidence of homology. The top hit was vCBM60 (PDB 2XHH; (45)), a family 60 carbohydrate-binding module from an uncultured bacterium, with a Z-score of 5.8 over 77 aligned residues at 3.4 Å root-mean-square deviation and 6% sequence identity.

## Data availability

All data supporting the conclusions of this study are included in the article and its supplemental material.

## ACKNOWLEDGEMENTS

The authors would like to thank members of the Bernhardt, Rudner, and Kruse labs for advice and helpful discussions. Piet de Boer kindly provided the anti-RodZ antiserum. Paula Montero Llopis and Microscopy Resources on the North Quad (MicRoN) at Harvard Medical School provided imaging training and support. This work was supported by the National Institute of Allergy and Infectious Diseases of the National Institutes of Health [(F31AI181426 to KAS) and (R01 AI083365 to TGB) and Investigator funds from the Howard Hughes Medical Institute (TGB). The content is solely the responsibility of the authors and does not necessarily represent the official views of the National Institutes of Health.

## SUPPLEMENTARY MATERIAL

**Fig. S1.**
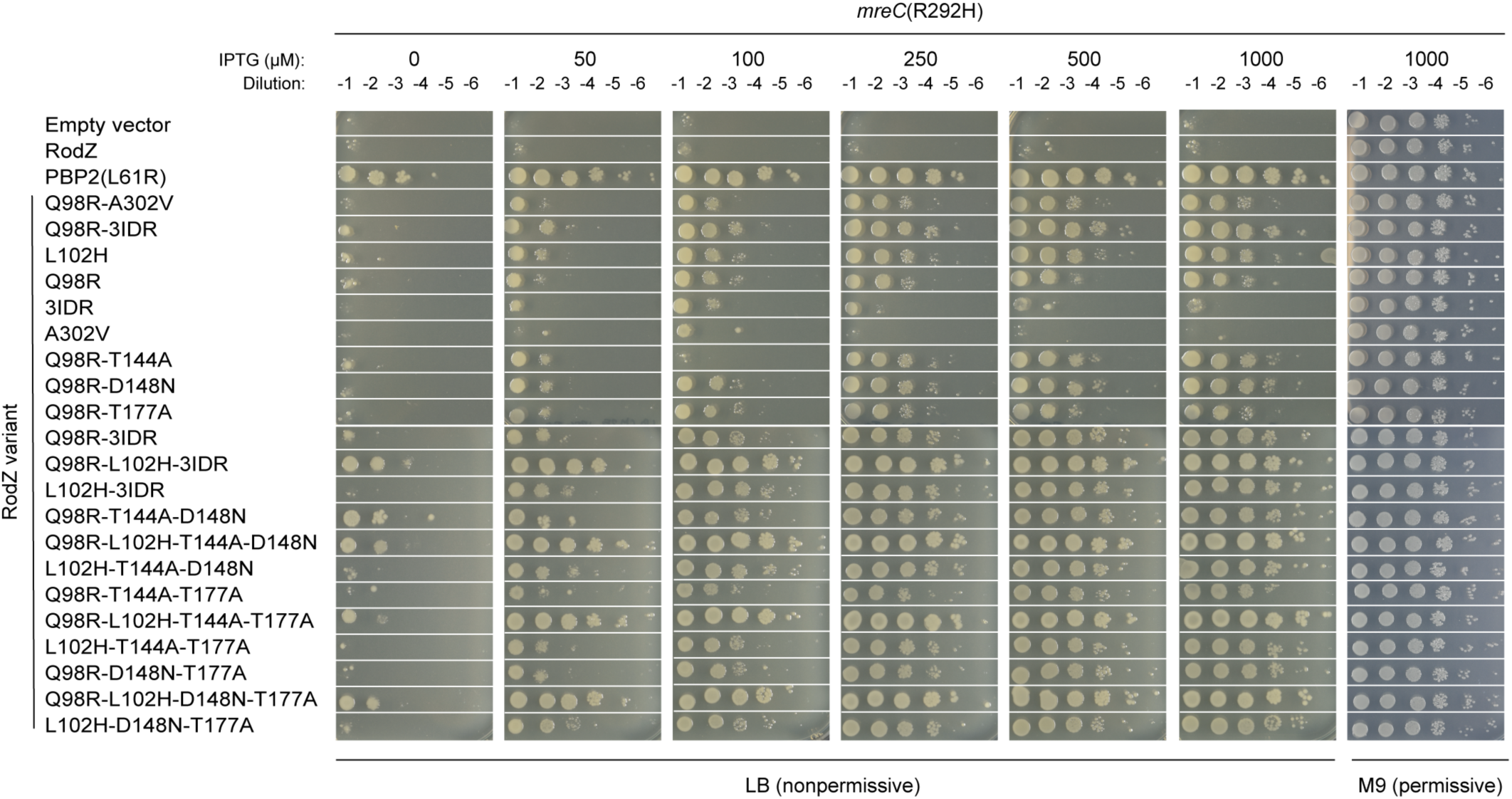
Testing additional RodZ variants for suppression of the *mreC(R292H)* growth defect. Spot-dilution viability assay of PR5 [*mreC(R292H)*] carrying the indicated P*_lac_*-driven *rodZ* alleles. Plates contained IPTG at 0, 50, 100, 250, 500, or 1,000 µM (LB) or 1,000 µM (M9 CAA glu) as indicated. Assay was performed as in Figure 2.

**Fig. S2.**
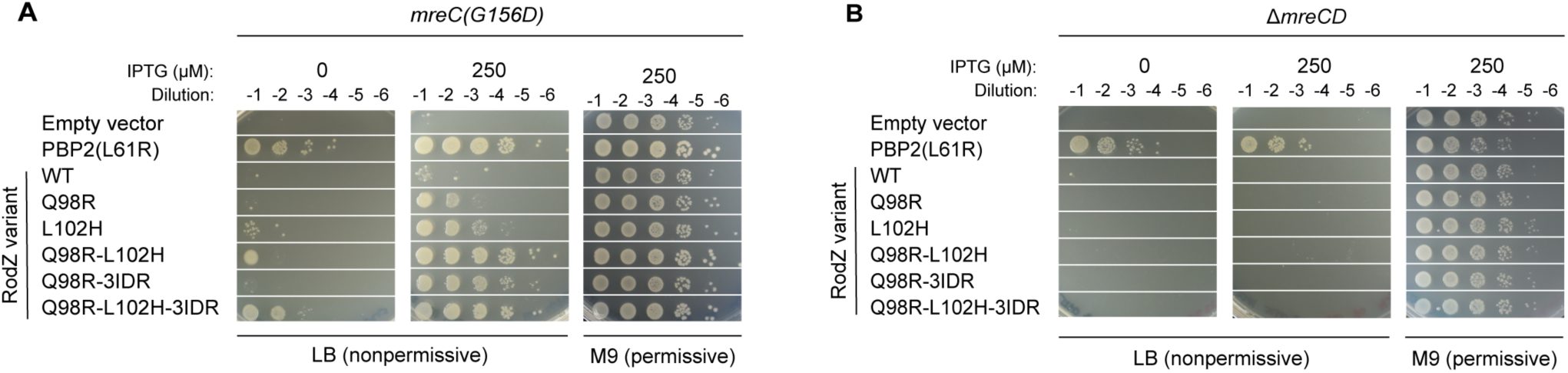
RodZ variants suppress *mreC*(G156D) but not Δ*mreCD* defects. Spot-dilution viability assays of PR30 [*mreC*(G156D)] **(A)** and PR137 [Δ*mreCD*] **(B)** strains expressing the indicated RodZ variants. Assays were performed as for Figure 2.

**Fig. S3.**
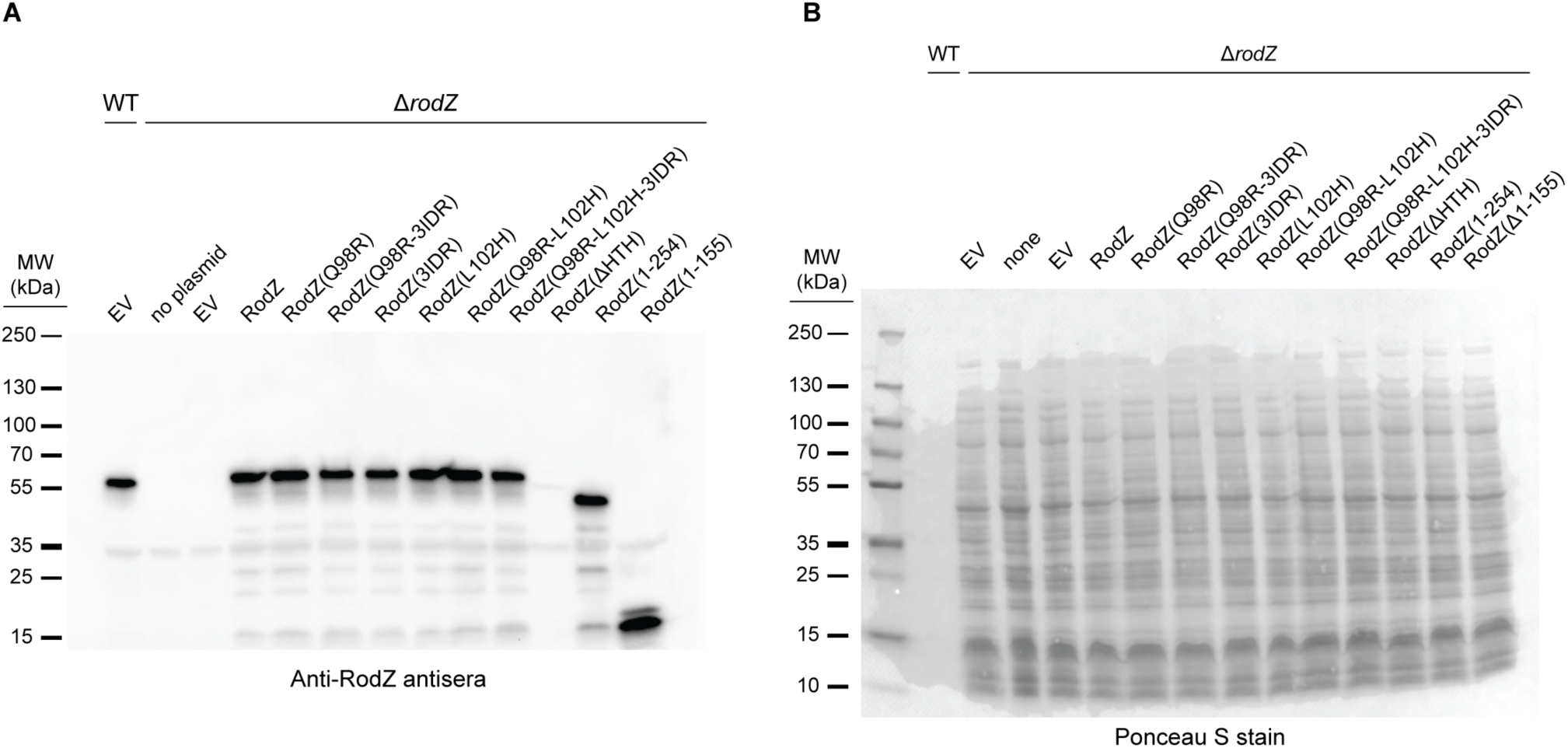
Immunoblot of RodZ variants. **(A)** Whole-cell extracts of WT and Δ*rodZ* cells producing the indicated RodZ variant were immunoblotted to detect RodZ using anti-RodZ antiserum. **(B)** The blot was also stained with Ponceau S stain to control for loading and blotting.

**Fig. S4.**
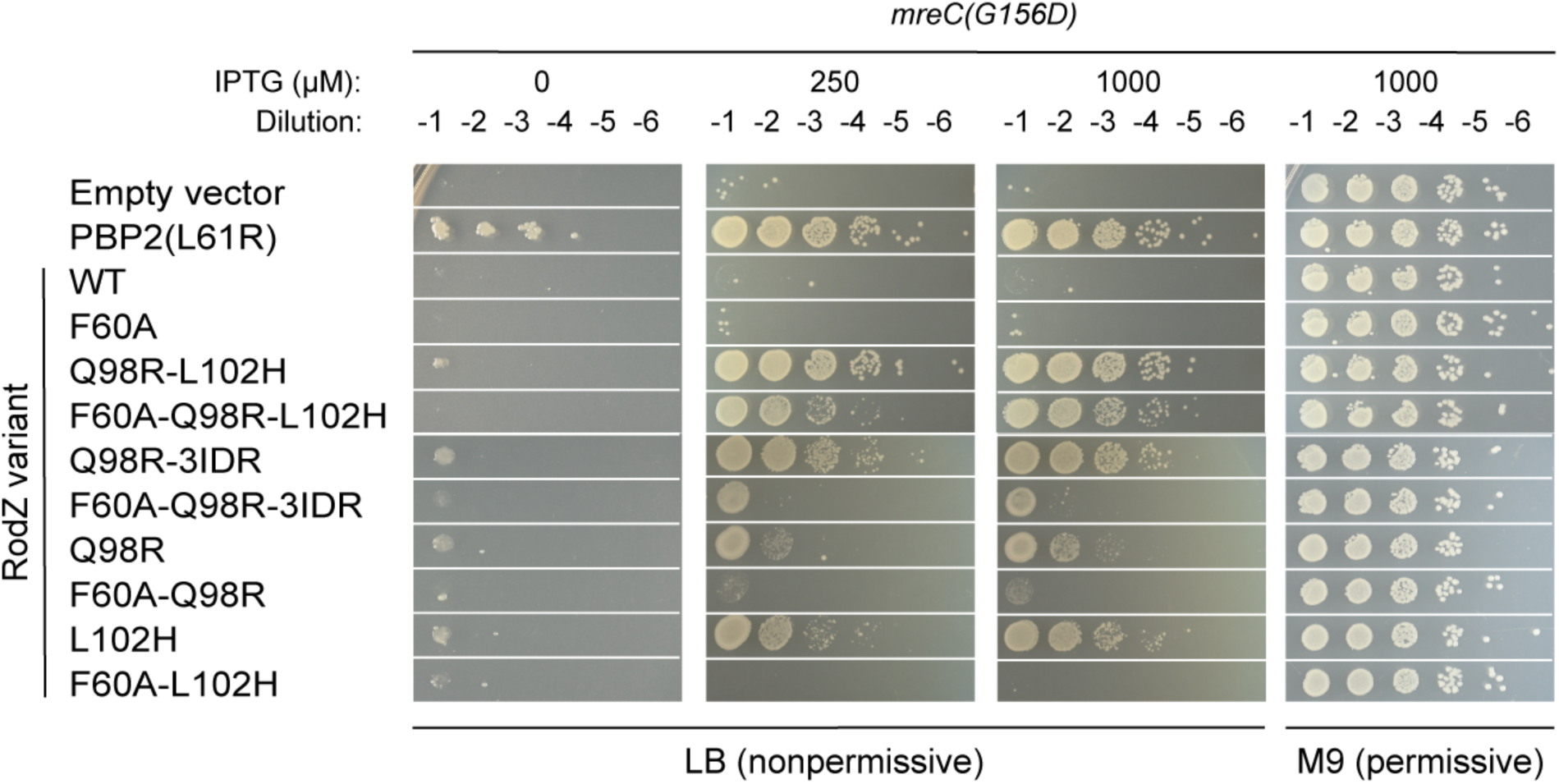
Affect of F60A substitution on the suppression of the *mreC*(G156D) growth defect. Spot-dilution viability assay of PR30 [*mreC*(G156D)] cells carrying the RodZ(F60A) variant series. Plates contained 1,000 µM IPTG (M9 CAA glu) or 0, 100, 250, or 1,000 µM IPTG (LB) as indicated. The assay was performed as described in Figure 2.

**Fig. S5.**
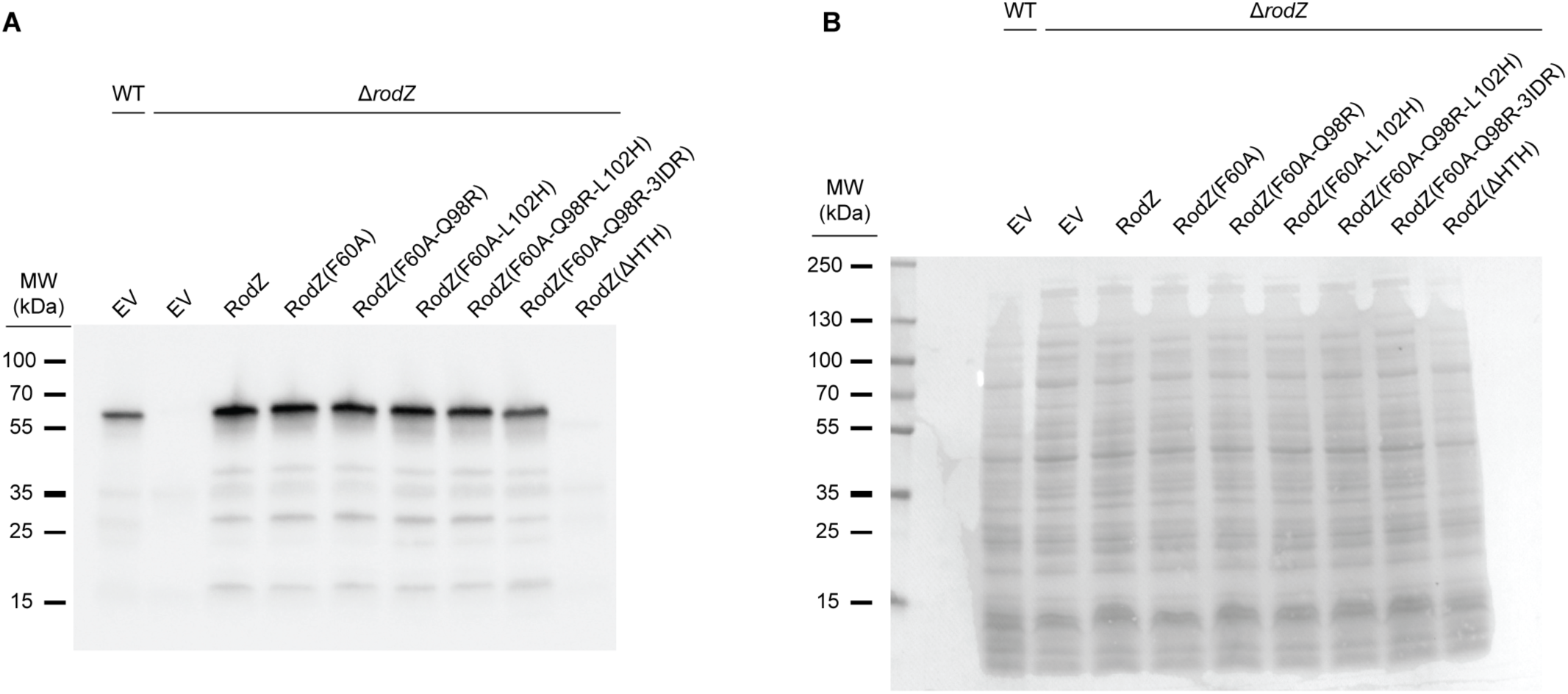
Immunoblot of RodZ(F60A) variants. **(A)** Whole-cell extracts of WT and Δ*rodZ* cells producing the indicated RodZ variant were immunoblotted to detect RodZ using anti-RodZ antiserum. **(B)** The blot was also stained with Ponceau S stain to control for loading and blotting.

**Table S1.** *rodZ* mutations identified in isolates and their corresponding RodZ domain.

| **Isolate #** | ***rodZ* mutation** | **RodZ protein change** | **Domain** |
|---|---|---|---|
| A13 | CAG → CGG | Q98R | JM |
|  | GCC → GTC | A302V | CFD |
| B3 | CTC → CAC | L102H | JM |
| B9 | CAG → CGG | Q98R | JM |
|  | ACG → GCG | T144A | IDR |
|  | GAT → AAT | D148N | IDR |
|  | ACG → GCG | T177A | IDR |

**Table S2.**
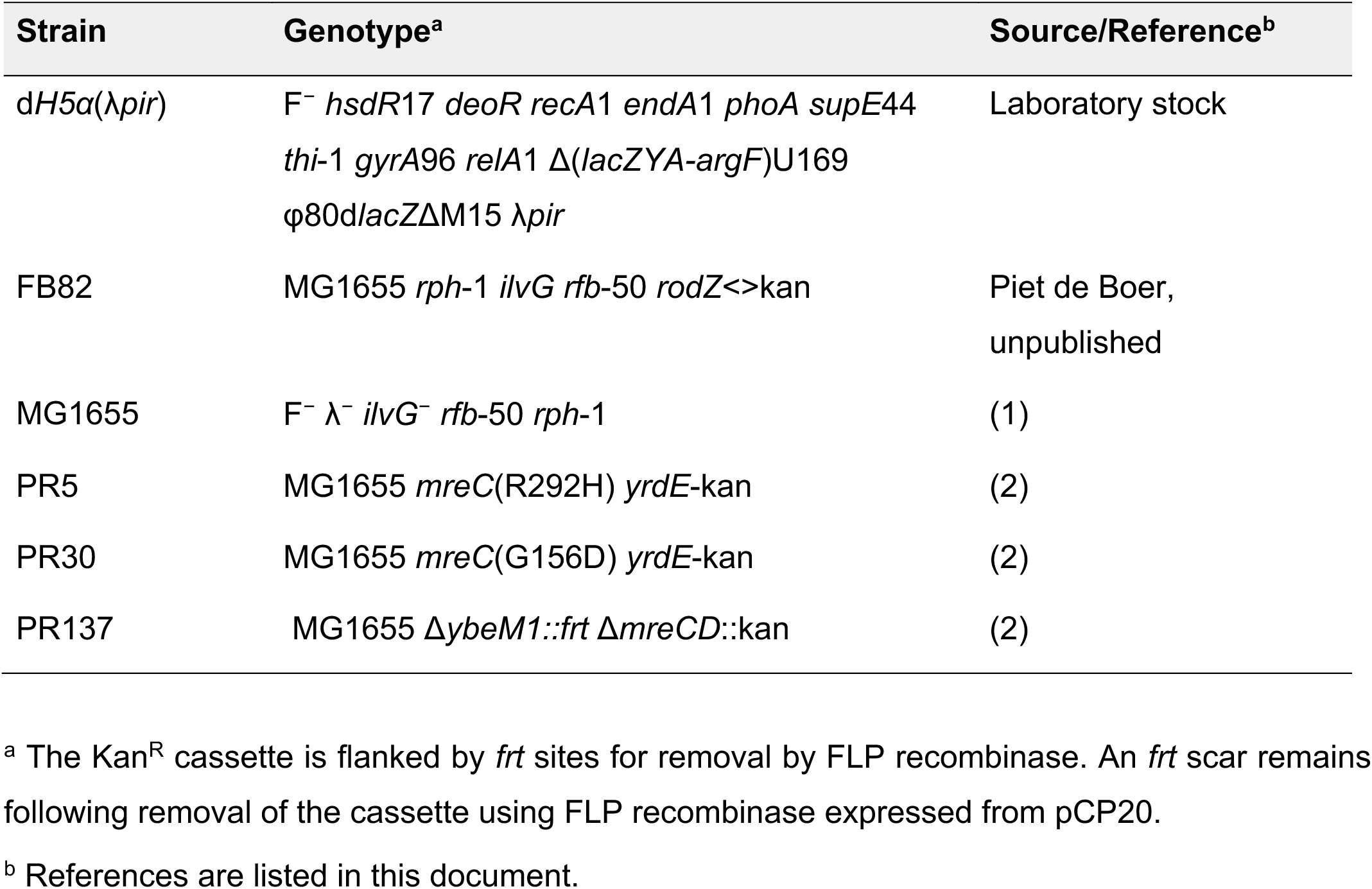
Strains used in this study.

| **Strain** | **Genotype<sup>a</sup>** | **Source/Reference<sup>b</sup>** |
|---|---|---|
| d*H5α*(λ*pir*) | F<sup>−</sup> *hsdR*17 *deoR* *recA*1 *endA*1 *phoA* *supE*44 *thi*-1 *gyrA*96 *relA*1 Δ(*lacZYA-argF*)U169 φ80d*lacZ*ΔM15 λ*pir* | Laboratory stock |
| FB82 | MG1655 *rph*-1 *ilvG* *rfb*-50 *rodZ*<>kan | Piet de Boer, unpublished |
| MG1655 | F<sup>−</sup> λ<sup>−</sup> *ilvG*<sup>−</sup> *rfb*-50 *rph*-1 | (1) |
| PR5 | MG1655 *mreC*(R292H) *yrdE*-kan | (2) |
| PR30 | MG1655 *mreC*(G156D) *yrdE*-kan | (2) |
| PR137 | MG1655 Δ*ybeM1::frt* Δ*mreCD*::kan | (2) |

<sup>a</sup> The Kan<sup>R</sup> cassette is flanked by *frt* sites for removal by FLP recombinase. An *frt* scar remains following removal of the cassette using FLP recombinase expressed from pCP20.
<sup>b</sup> References are listed in this document.

**Table S3.** Plasmids used in this study.

| Plasmid | Genotype<sup>a</sup> | Origin | Source or Reference<sup>b</sup> |
|---|---|---|---|
| pKS12 | *cat lacI<sup>q</sup>* P<sub>lac</sub>::strongRBS-*rodZ* | pACYC/p15a | This study |
| pKS13 | *cat lacI<sup>q</sup>* P<sub>lac</sub>::strongRBS-*rodZ*(1-155) | pACYC/p15a | This study |
| pKS14 | *cat lacI<sup>q</sup>* P<sub>lac</sub>::strongRBS-*rodZ*(85-337) | pACYC/p15a | This study |
| pKS28 | *cat lacI<sup>q</sup>* P<sub>lac</sub>::strongRBS-*rodZ*(1-254) | pACYC/p15a | This study |
| pKS49 | *cat lacI<sup>q</sup>* P<sub>lac</sub>::strongRBS-*rodZ*(Q98R) | pACYC/p15a | This study |
| pKS52 | *cat lacI<sup>q</sup>* P<sub>lac</sub>::strongRBS-*rodZ*(T177A) | pACYC/p15a | This study |
| pKS53 | *cat lacI<sup>q</sup>* P<sub>lac</sub>::strongRBS-*rodZ*(L102H) | pACYC/p15a | This study |
| pKS54 | *cat lacI<sup>q</sup>* P<sub>lac</sub>::strongRBS-*rodZ*(A302V) | pACYC/p15a | This study |
| pKS61 | *cat lacI<sup>q</sup>* P<sub>lac</sub>::strongRBS-*rodZ*(Q98R-L102H) | pACYC/p15a | This study |
| pKS63 | *cat lacI<sup>q</sup>* P<sub>lac</sub>::strongRBS-*rodZ*(1-155, Q98R) | pACYC/p15a | This study |
| pKS64 | *cat lacI<sup>q</sup>* P<sub>lac</sub>::strongRBS-*rodZ*(1-155, L102H) | pACYC/p15a | This study |
| pKS65 | *cat lacI<sup>q</sup>* P<sub>lac</sub>::strongRBS-*rodZ*(1-155, Q98R-L102H) | pACYC/p15a | This study |
| pKS66 | *cat lacI<sup>q</sup>* P<sub>lac</sub>::strongRBS-*rodZ*(1-254, Q98R) | pACYC/p15a | This study |
| pKS67 | *cat lacI<sup>q</sup>* P<sub>lac</sub>::strongRBS-*rodZ*(1-254, L102H) | pACYC/p15a | This study |
| pKS68 | *cat lacI<sup>q</sup>* P<sub>lac</sub>::strongRBS-*rodZ*(1-254, Q98R-L102H) | pACYC/p15a | This study |
| pKS69 | *cat lacI<sup>q</sup>* P<sub>lac</sub>::strongRBS-*rodZ*(Q98R-T144A) | pACYC/p15a | This study |
| pKS70 | *cat lacI<sup>q</sup>* P<sub>lac</sub>::strongRBS-*rodZ*(Q98R-D148N) | pACYC/p15a | This study |
| pKS71 | *cat lacI<sup>q</sup>* P<sub>lac</sub>::strongRBS-*rodZ*(Q98R-T177A) | pACYC/p15a | This study |
| pKS72 | *cat lacI<sup>q</sup>* P<sub>lac</sub>::strongRBS-*rodZ*(T144A-D148N) | pACYC/p15a | This study |
| pKS73 | *cat lacI<sup>q</sup>* P<sub>lac</sub>::strongRBS-*rodZ*(T144A-T177A) | pACYC/p15a | This study |
| pKS74 | *cat lacI<sup>q</sup>* P<sub>lac</sub>::strongRBS-*rodZ*(D148N-T177A) | pACYC/p15a | This study |
| pKS76 | *cat lacI<sup>q</sup>* P<sub>lac</sub>::strongRBS-*rodZ*(Q98R-T144A-D148N) | pACYC/p15a | This study |
| pKS77 | *cat lacI<sup>q</sup>* P<sub>lac</sub>::strongRBS-*rodZ*(Q98R-L102H-T144A-D148N) | pACYC/p15a | This study |
| pKS78 | *cat lacI<sup>q</sup>* P<sub>lac</sub>::strongRBS-*rodZ*(L102H-T144A-D148N) | pACYC/p15a | This study |
| pKS79 | *cat lacI<sup>q</sup>* P<sub>lac</sub>::strongRBS-*rodZ*(Q98R-T144A-T177A) | pACYC/p15a | This study |
| pKS80 | *cat lacI<sup>q</sup>* P<sub>lac</sub>::strongRBS-*rodZ*(Q98R-L102H-T144A-T177A) | pACYC/p15a | This study |
| pKS81 | *cat lacI<sup>q</sup>* P<sub>lac</sub>::strongRBS-*rodZ*(L102H-T144A-T177A) | pACYC/p15a | This study |
| pKS82 | *cat lacI<sup>q</sup>* P<sub>lac</sub>::strongRBS-*rodZ*(Q98R-D148N-T177A) | pACYC/p15a | This study |
| pKS83 | *cat lacI<sup>q</sup>* P<sub>lac</sub>::strongRBS-*rodZ*(Q98R-L102H-D148N-T177A) | pACYC/p15a | This study |
| pKS84 | *cat lacI<sup>q</sup>* P<sub>lac</sub>::strongRBS-*rodZ*(L102H-D148N-T177A) | pACYC/p15a | This study |
| pKS89 | *cat lacI<sup>q</sup>* P<sub>lac</sub>::strongRBS-*rodZ*(Q98R-A302V) | pACYC/p15a | This study |
| pKS90 | *cat lacI<sup>q</sup>* P<sub>lac</sub>::strongRBS-*rodZ*(T144A-D148N-T177A) | pACYC/p15a | This study |
| pKS91 | *cat lacI<sup>q</sup>* P<sub>lac</sub>::strongRBS-*rodZ*(Q98R-T144A-D148N-T177A) | pACYC/p15a | This study |
| pKS92 | *cat lacI<sup>q</sup>* P<sub>lac</sub>::strongRBS-*rodZ*(Q98R-L102H-T144A-D148N-T177A) | pACYC/p15a | This study |
| pKS93 | *cat lacI<sup>q</sup>* P<sub>lac</sub>::strongRBS-*rodZ*(L102H-T144A-D148N-T177A) | pACYC/p15a | This study |
| pKS116 | *cat lacI<sup>q</sup>* P<sub>lac</sub>::strongRBS-*rodZ*(F60A) | pACYC/p15a | This study |
| pKS117 | *cat lacI<sup>q</sup>* P<sub>lac</sub>::strongRBS-*rodZ*(F60A-Q98R-L102H) | pACYC/p15a | This study |
| pKS118 | *cat lacI<sup>q</sup>* P<sub>lac</sub>::strongRBS-*rodZ*(F60A-Q98R-T144A-D148N-T177A) | pACYC/p15a | This study |
| pKS119 | *cat lacI<sup>q</sup>* P<sub>lac</sub>::strongRBS-*rodZ*(F60A-Q98R) | pACYC/p15a | This study |
| pKS120 | *cat lacI<sup>q</sup>* P<sub>lac</sub>::strongRBS-*rodZ*(F60A-L102H) | pACYC/p15a | This study |
| pMT77 | *cat lacI<sup>q</sup>* P<sub>lac</sub>::empty | pACYC/p15a | Lab stock |
| pPR122 | *cat* P<sub>lac</sub>::nativeRBS-*pbpA*(L61R)-*rodA* | pBR/ColE1 | (3) |

<sup>a</sup> Gene fusions and point substitutions are indicated in the genotype column.
<sup>b</sup> References are listed in this document.

**Table S4.**
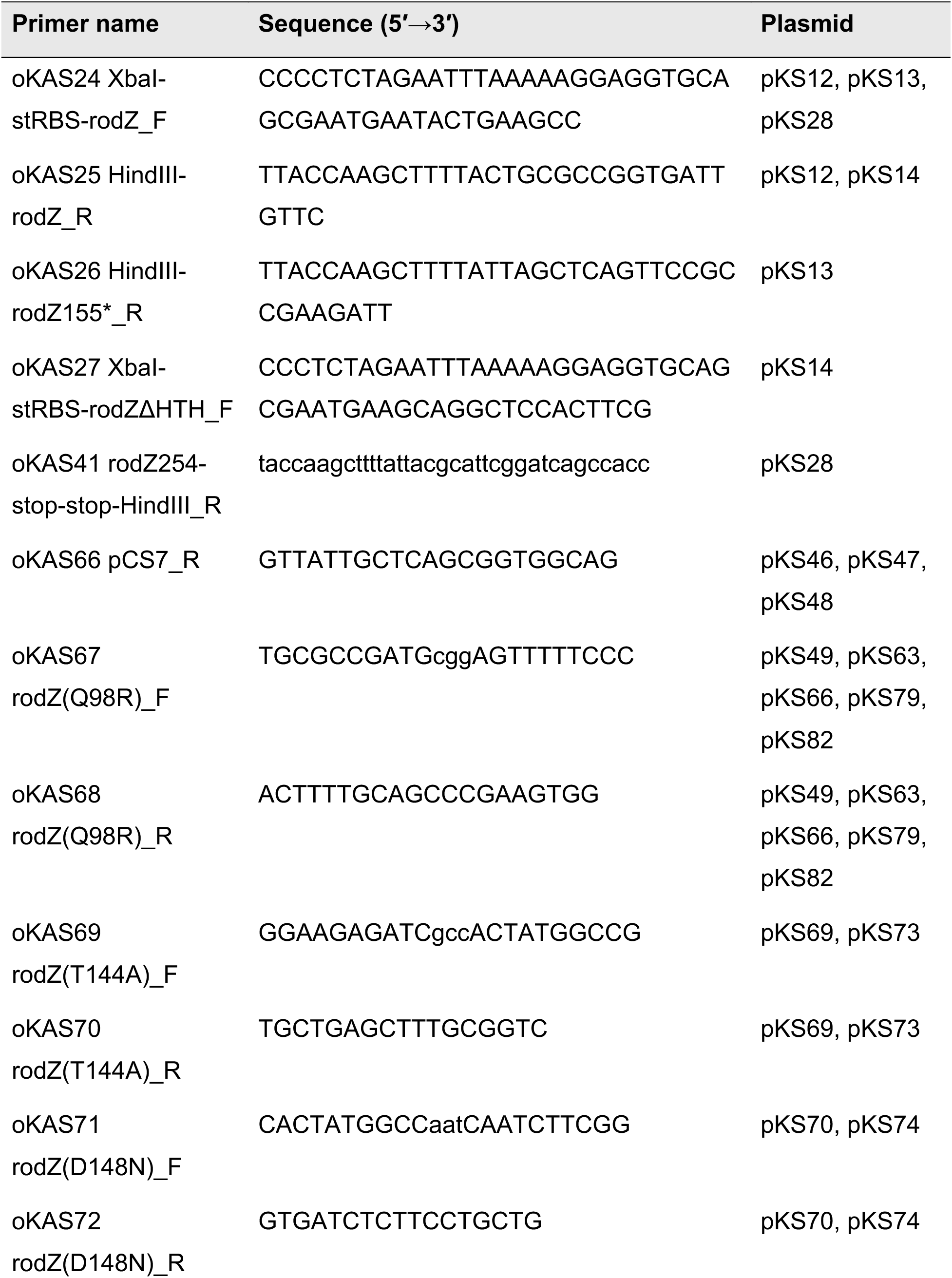
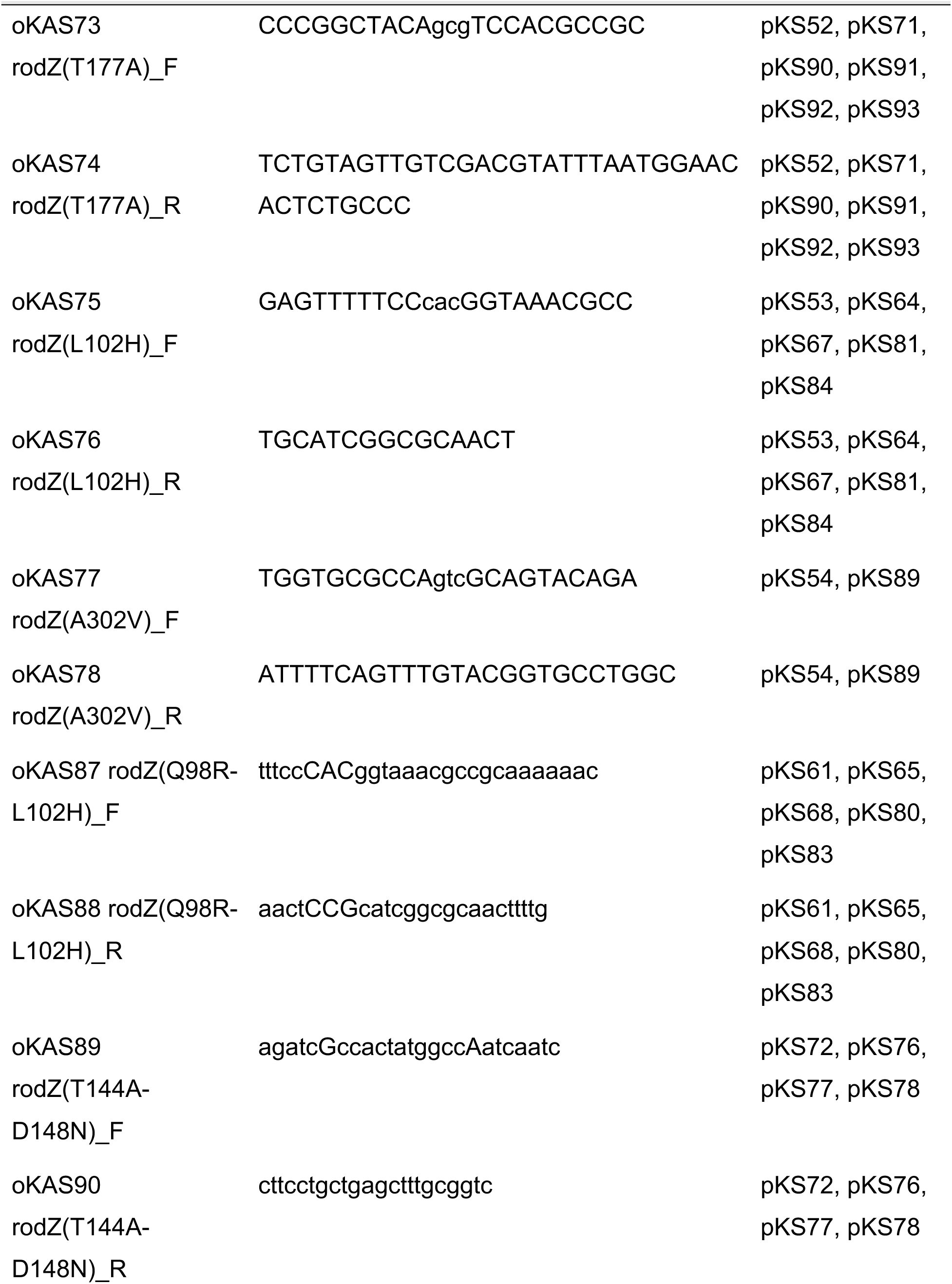
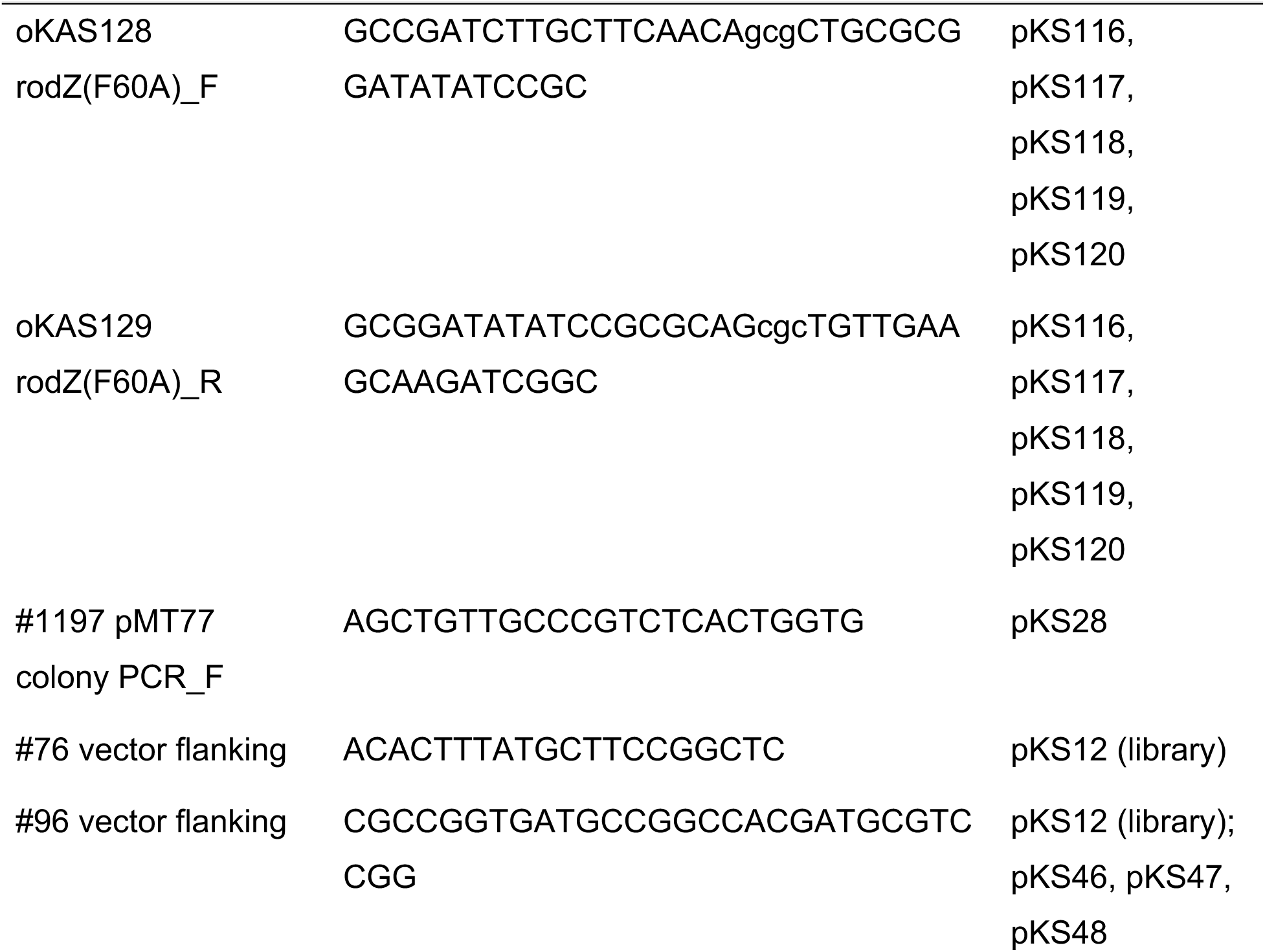
Primers used in this study.

## SUPLEMENTAL METHODS

### Plasmid Construction

All constructs were verified by Sanger sequencing or whole-plasmid sequencing. Point mutations were introduced by KLD mutagenesis (kinase-ligase-DpnI treatment of a whole-plasmid PCR product, ∼4.4 kb).

### pKS12

The *rodZ* insert was amplified from MG1655 genomic DNA using primers oKAS24 (CCCCTCTAGAATTTAAAAAGGAGGTGCAGCGAATGAATACTGAAGCC) and oKAS25 (TTACCAAGCTTTTACTGCGCCGGTGATTGTTC), and the 1,057-bp product was inserted into pMT77 using restriction enzymes XbaI and HindIII. The native ribosome-binding site was replaced with a strong ribosome-binding site (AGGAGG) encoded in oKAS24.

### pKS13

The *rodZ*(1–155) insert was amplified from MG1655 genomic DNA using primers oKAS24 (CCCCTCTAGAATTTAAAAAGGAGGTGCAGCGAATGAATACTGAAGCC) and oKAS26 (TTACCAAGCTTTTATTAGCTCAGTTCCGCCGAAGATT), and the 514-bp product was inserted into pMT77 using restriction enzymes XbaI and HindIII.

### pKS14

The *rodZ*(85–337) insert was amplified from MG1655 genomic DNA using primers oKAS27 (CCCTCTAGAATTTAAAAAGGAGGTGCAGCGAATGAAGCAGGCTCCACTTCG) and oKAS25 (TTACCAAGCTTTTACTGCGCCGGTGATTGTTC), and the 807-bp product was inserted into pMT77 using restriction enzymes XbaI and HindIII.

### pKS28

The *rodZ*(1–254) insert was amplified from MG1655 genomic DNA using primers oKAS24 (CCCCTCTAGAATTTAAAAAGGAGGTGCAGCGAATGAATACTGAAGCC) and oKAS41 (taccaagcttttattacgcattcggatcagccacc), and the 810-bp product was inserted into pMT77 using restriction enzymes XbaI and HindIII. Transformants were screened by colony PCR using primers #1197 (AGCTGTTGCCCGTCTCACTGGTG) and oKAS41, which yield a 1,152-bp product from the correct construct.

### pKS49

Generated by KLD mutagenesis using pKS12 as a template and primers oKAS67 (TGCGCCGATGcggAGTTTTTCCC) and oKAS68 (ACTTTTGCAGCCCGAAGTGG).

### pKS52

Generated by KLD mutagenesis using pKS12 as a template and primers oKAS73 (CCCGGCTACAgcgTCCACGCCGC) and oKAS74 (TCTGTAGTTGTCGACGTATTTAATGGAACACTCTGCCC).

### pKS53

Generated by KLD mutagenesis using pKS12 as a template and primers oKAS75 (GAGTTTTTCCcacGGTAAACGCC) and oKAS76 (TGCATCGGCGCAACT).

### pKS54

Generated by KLD mutagenesis using pKS12 as a template and primers oKAS77 (TGGTGCGCCAgtcGCAGTACAGA) and oKAS78 (ATTTTCAGTTTGTACGGTGCCTGGC).

### pKS61

Generated by KLD mutagenesis using pKS12 as a template and primers oKAS87 (tttccCACggtaaacgccgcaaaaaac) and oKAS88 (aactCCGcatcggcgcaacttttg). The primer pair introduces both the Q98R and L102H substitutions in a single reaction.

### pKS63

Generated by KLD mutagenesis using pKS13 as a template and primers oKAS67 (TGCGCCGATGcggAGTTTTTCCC) and oKAS68 (ACTTTTGCAGCCCGAAGTGG).

### pKS64

Generated by KLD mutagenesis using pKS13 as a template and (GAGTTTTTCCcacGGTAAACGCC) and oKAS76 (TGCATCGGCGCAACT). primers oKAS75

### pKS65

Generated by KLD mutagenesis using pKS13 as a template and primers oKAS87 (tttccCACggtaaacgccgcaaaaaac) and oKAS88 (aactCCGcatcggcgcaacttttg).

### pKS66

Generated by KLD mutagenesis using pKS28 as a template and primers oKAS67 (TGCGCCGATGcggAGTTTTTCCC) and oKAS68 (ACTTTTGCAGCCCGAAGTGG).

### pKS67

Generated by KLD mutagenesis using pKS28 as a template and primers oKAS75 (GAGTTTTTCCcacGGTAAACGCC) and oKAS76 (TGCATCGGCGCAACT).

### pKS68

Generated by KLD mutagenesis using pKS28 as a template and primers oKAS87 (tttccCACggtaaacgccgcaaaaaac) and oKAS88 (aactCCGcatcggcgcaacttttg).

### pKS69

Generated by KLD mutagenesis using pKS49 as a template and primers oKAS69 (GGAAGAGATCgccACTATGGCCG) and oKAS70 (TGCTGAGCTTTGCGGTC).

### pKS70

Generated by KLD mutagenesis using pKS49 as a template and primers oKAS71 (CACTATGGCCaatCAATCTTCGG) and oKAS72 (GTGATCTCTTCCTGCTG).

### pKS71

Generated by KLD mutagenesis using pKS49 as a template and primers oKAS73 (CCCGGCTACAgcgTCCACGCCGC) and oKAS74 (TCTGTAGTTGTCGACGTATTTAATGGAACACTCTGCCC).

### pKS72

Generated by KLD mutagenesis using pKS12 as a template and primers oKAS89 (agatcGccactatggccAatcaatc) and oKAS90 (cttcctgctgagctttgcggtc).

### pKS73

Generated by KLD mutagenesis using pKS52 as a template and primers oKAS69 (GGAAGAGATCgccACTATGGCCG) and oKAS70 (TGCTGAGCTTTGCGGTC).

### pKS74

Generated by KLD mutagenesis using pKS52 as a template and primers oKAS71 (CACTATGGCCaatCAATCTTCGG) and oKAS72 (GTGATCTCTTCCTGCTG). Annealing temperature 60°C.

### pKS76

Generated by KLD mutagenesis using pKS49 as a template and primers oKAS89 (agatcGccactatggccAatcaatc) and oKAS90 (cttcctgctgagctttgcggtc). The primer pair introduces both the T144A and D148N substitutions in a single reaction.

### pKS77

Generated by KLD mutagenesis using pKS61 as a template and primers oKAS89 (agatcGccactatggccAatcaatc) and oKAS90 (cttcctgctgagctttgcggtc).

### pKS78

Generated by KLD mutagenesis using pKS53 as a template and primers oKAS89 (agatcGccactatggccAatcaatc) and oKAS90 (cttcctgctgagctttgcggtc).

### pKS79

Generated by KLD mutagenesis using pKS73 as a template and primers oKAS67 (TGCGCCGATGcggAGTTTTTCCC) and oKAS68 (ACTTTTGCAGCCCGAAGTGG).

### pKS80

Generated by KLD mutagenesis using pKS73 as a template and primers oKAS87 (tttccCACggtaaacgccgcaaaaaac) and oKAS88 (aactCCGcatcggcgcaacttttg).

### pKS81

Generated by KLD mutagenesis using pKS73 as a template and primers oKAS75 (GAGTTTTTCCcacGGTAAACGCC) and oKAS76 (TGCATCGGCGCAACT).

### pKS82

Generated by KLD mutagenesis using pKS74 as a template and primers oKAS67 (TGCGCCGATGcggAGTTTTTCCC) and oKAS68 (ACTTTTGCAGCCCGAAGTGG).

### pKS83

Generated by KLD mutagenesis using pKS74 as a template and primers oKAS87 (tttccCACggtaaacgccgcaaaaaac) and oKAS88 (aactCCGcatcggcgcaacttttg).

### pKS84

Generated by KLD mutagenesis using pKS74 as a template and primers oKAS75 (GAGTTTTTCCcacGGTAAACGCC) and oKAS76 (TGCATCGGCGCAACT).

### pKS89

Generated by KLD mutagenesis using pKS49 as a template and primers oKAS77 (TGGTGCGCCAgtcGCAGTACAGA) and oKAS78 (ATTTTCAGTTTGTACGGTGCCTGGC).

### pKS90

Generated by KLD mutagenesis using pKS72 as a template and primers oKAS73 (CCCGGCTACAgcgTCCACGCCGC) and oKAS74 (TCTGTAGTTGTCGACGTATTTAATGGAACACTCTGCCC).

### pKS91

Generated by KLD mutagenesis using pKS76 as a template and primers oKAS73 (CCCGGCTACAgcgTCCACGCCGC) and oKAS74 (TCTGTAGTTGTCGACGTATTTAATGGAACACTCTGCCC).

### pKS92

Generated by KLD mutagenesis using pKS77 as a template and primers oKAS73 (CCCGGCTACAgcgTCCACGCCGC) and oKAS74 (TCTGTAGTTGTCGACGTATTTAATGGAACACTCTGCCC).

### pKS93

Generated by KLD mutagenesis using pKS78 as a template and primers oKAS73 (CCCGGCTACAgcgTCCACGCCGC) and oKAS74 (TCTGTAGTTGTCGACGTATTTAATGGAACACTCTGCCC).

### pKS116

Generated by KLD mutagenesis using pKS12 as a template and primers oKAS128 (GCCGATCTTGCTTCAACAgcgCTGCGCGGATATATCCGC) and oKAS129 (GCGGATATATCCGCGCAGcgcTGTTGAAGCAAGATCGGC).

### pKS117

Generated by KLD mutagenesis using pKS61 as a template and primers oKAS128 (GCCGATCTTGCTTCAACAgcgCTGCGCGGATATATCCGC) and oKAS129 (GCGGATATATCCGCGCAGcgcTGTTGAAGCAAGATCGGC).

### pKS118

Generated by KLD mutagenesis using pKS91 as a template and primers oKAS128 (GCCGATCTTGCTTCAACAgcgCTGCGCGGATATATCCGC) and oKAS129 (GCGGATATATCCGCGCAGcgcTGTTGAAGCAAGATCGGC).

### pKS119

Generated by KLD mutagenesis using pKS49 as a template (GCCGATCTTGCTTCAACAgcgCTGCGCGGATATATCCGC) (GCGGATATATCCGCGCAGcgcTGTTGAAGCAAGATCGGC). and and primers oKAS128 oKAS129

### pKS120

Generated by KLD mutagenesis using pKS53 as a template (GCCGATCTTGCTTCAACAgcgCTGCGCGGATATATCCGC) and primers oKAS128 and oKAS129 (GCGGATATATCCGCGCAGcgcTGTTGAAGCAAGATCGGC).

